# Computational Support, Not Primacy, Distinguishes Compensatory Memory Reorganization in Epilepsy

**DOI:** 10.1101/2020.06.07.138594

**Authors:** Joseph I. Tracy, Kapil Chaudhary, Shilpi Modi, Andrew Crow, Ashith Kumar, David Weinstein, Michael R. Sperling

## Abstract

Temporal lobe epilepsy (TLE) is associated with impairment in episodic memory. A substantial sub-group, however, is able to maintain adequate memory despite temporal lobe pathology. What has been missing from prior work in the cognitive reorganization is a direct comparison of TLE patients with intact/compensated status from those who are memory impaired and uncompensated. Little is known about the particular regional activations, functional connectivities (FC’s), and/or network reconfigurations that implement changes in the primary computations or the support functions that drive adaptive plasticity and compensated memory. We utilized task fMRI on 54 unilateral TLE patients and 24 matched healthy controls (HC) during performance of a paired-associate memory (PAM) task to address three questions: 1) what regions implement PAM in TLE, and do such regions vary as a function of good versus poor performance, 2) are there unique FC’s present during memory encoding that accounts for intact status through the preservation of primary memory computations or the supportive computations that allow for compensated memory responses, and 3) what features during memory encoding are most distinctive: is it the magnitude and location of regional activations, or the presence of enhanced functional connections to key structures such as the hippocampus? Results revealed a unique profile of non-ictal, non-dominant hemisphere regions (e.g., right posterior temporal regions) were most important to intact/compensated status in LTLE, involving both increased regional activity and increased modulatory communication with the hippocampi, all feature that was missing in impaired/uncompensated LTLE. The profile involved areas that are neither contralateral homologues to left hemisphere memory areas, nor regions traditionally considered computationally primary for episodic memory. None of these areas of increased activation or functional connectivity were associated with advantaged memory in HC’s. Our emphasis on different performance levels yielded insight into two forms of cognitive reorganization. Computational primacy, where LTLE showed little change relative to HC’s, and computational support where Intact/Compensated LTLE patients showed adaptive abnormalities. The analyses isolated the unique regional activations and mediating FC’s that implement truly compensatory reorganization in LTLE. The results provided a new perspective on memory deficits by making clear that memory deficits arise not just from knockout of a functional hub, but from the failure to instantiate a complex set of reorganization responses. Such responses provided computational support to ensure successful memory. The findings demonstrated that by keeping track of performance levels, we can increase our understanding of adaptive brain responses and neuroplasticity in epilepsy.

## Introduction

Temporal lobe epilepsy (TLE) is associated with cognitive impairment, most commonly in episodic memory, with up to 70% of TLE patients displaying memory problems (Gleissner *et al.*, 2002; Flugel *et al.*, 2006; Glikmann-Johnston *et al.*, 2008; Saling, 2009). There is a substantial sub-group, however, who are able to maintain adequate episodic memory despite their temporal lobe disease (Coras *et al.*, 2014). While the functional neuroanatomy of episodic memory deficits has been studied and explicated, we know very little about the regional and brain network features that support preserved memory in the setting of temporal lobe disease. The subcortical region, the hippocampus, plays a major role in seizure generation and spread in TLE (McIntyre and Racine, 1986), and mesial temporal sclerosis involving hippocampal atrophy remains the most common pathology of focal TLE (Liu *et al.*, 1995). The hippocampus is also critically important to the formation of episodic long-term memories, through its known specialization for associative encoding and memory consolidation emerging from animal models (Squire and Zola-Morgan, 1991), electrophysiology (Burke *et al.*, 2014*a*,*b*, 2015; Long and Kahana, 2015; Norman, 2010; Yassa and Stark, 2011) and task-fMRI. For instance, task-based functional MRI (fMRI) studies with healthy individuals have demonstrated verbal memory tasks reliably activate the medial temporal lobe including hippocampus and para-hippocampal regions, particularly when evoked by associative encoding such as paired-associate learning paradigms (Clark *et al.*, 2018; Gould *et al.*, 2003; Law *et al.*, 2005; Meltzer and Constable, 2005; Mottaghy *et al.*, 1999; Vannest *et al.*, 2012). Accordingly, impairments in learning and memory are common in TLE, and removal of the hippocampus from surgery to control seizures often has a negative effect on episodic memory (Baxendale *et al*., 2006; Bowles *et al*., 2010; Helmstaedter and Elger, 1996; Khalil *et al*., 2016; Manns and Eichenbaum, 2006; Mueller *et al*., 2012).

In the setting of temporal lobe pathology, the brain’s adaptive drive to maintain adequate memory performance can result in cognitive reorganization, involving the redistribution both of primary memory computations and alterations in the use and recruitment of the regions and networks that provide the supportive processing, i.e., the necessary but not sufficient functions to implement effective memory. Several previous studies have shown intra- and inter-hemispheric reorganization in memory encoding networks across both verbal and visual domains in individuals with left and right TLE (Alessio *et al.*, 2013; Bell and Davies, 1998; Bonelli *et al.*, 2010; Sidhu *et al.*, 2013; Kim *et al.*, 2003; Richardson *et al.*, 2003, 2006; Milian *et al.*, 2015; Powell *et al.*, 2007). In TLE, paired-associate memory tasks (Milian *et al.*, 2015; Smith *et al.*, 2011) have been quite effective at revealing the reorganization of memory-related networks, involving structures such as the mesial temporal lobe and hippocampus. The presumptive goal of such reorganization is to compensate for the dysfunctional epileptogenic region. There has been prior work describing the general patterns of cognitive reorganization that might help maintain or restore adaptive functioning in the setting of brain disease such as temporal lobe epilepsy (Tracy and Osipowicz, 2011). What has been missing from the above studies, however, is a direct comparison of TLE patients with intact, compensated memory functioning from those who are memory impaired and uncompensated. The degree to which TLE patients with intact memory actually differ from normal controls with similar levels of memory performance also remains unchartered and poorly understood. More specifically, in the setting of temporal lobe disease, little is known about the particular regional activations, functional connectivities, and/or network reconfigurations that appear distinctly prone to implement either changes in the primary computations or the supportive functions that drive adaptive plasticity and compensated memory output (Bonelli *et al.*, 2013; Cheung *et al.*, 2009; Limotai *et al.*, 2018; Sidhu *et al.*, 2016; Tracy and Doucet, 2015).

With these advances in mind, we examined regional activation and network connectivities in TLE, but do so with a focus on important subgroups, namely, those whose memory performance suggests either intact/compensated or impaired/uncompensated brain responses. Utilizing task fMRI, we addressed three questions: First, what regions implement paired-associate memory (PAM) in TLE compared to healthy controls, and do such regions vary as a function of good versus poor PAM performance? Second, are there unique functional connectivities present during memory encoding that may account for intact status in TLE through the preservation of primary memory computations or the supportive functions that allow for cognitively compensated performance? Third, what are the features during memory encoding that are most distinctive about intact/compensated memory: is it the magnitude and location of regional activations during memory encoding, or the presence of enhanced functional connections to key structures such as the hippocampus?

We first tested for regional activation differences between our experimental groups, focusing on TLE group differences from healthy matched controls (HC). Next, after categorizing participants as good versus poor PAM performers, we utilized a two-factor GLM model to determine if such regional activations varied by Experimental Group (TLE, HC) and PAM performance, with the interaction of these factors the effect of key interest. To more carefully delineate cognitive compensation within our TLE patients, we separated TLE patients with intact and compensated performance from those with clearly impaired PAM performance. Based upon the above performance distinctions, we then utilized generalized psychophysiological interaction techniques (gPPI) to capture the differential functional connectivity present during successfully remembered PAM trials compared to a non-memory control condition. In these gPPI analyses, our focus was on whole brain connectivity to the region(s) producing key activation differences between our performance groups, i.e., the hippocampus, a region with well-established importance to episodic memory functioning. Lastly, we leveraged the classification power of Support Vector Machine Learning (SVM), to identify the features in our data that best discriminated intact/compensated versus impaired/uncompensated performance in TLE. More specifically, SVM determined whether the magnitude of regional effects sizes on the PAM task, as opposed to the functional connectivities mediating successful encoding, constituted better classifiers of intact/compensated status. The goal of this last analysis was to isolate the characteristics most robustly associated with truly successful verbal memory encoding and adaptive brain functional reorganization in TLE.

## Material and methods

A total of 54 patients with drug-resistant unilateral TLE (Left= 32; Right= 22) matched on age, handedness (Oldfield, 1971), and gender were recruited from the Thomas Jefferson University Comprehensive Epilepsy Centre. All patients were determined to be surgical candidates for either a standard anterior temporal lobectomy or thermal ablation of the ictal mesial temporal lobe based upon a multimodal evaluation including neurologic history and examination, scalp video-EEG, MRI, PET, and neuropsychological testing (Sperling *et al.*, 1996). All participants were left hemisphere language dominant, as verified by an fMRI verb generation task. All patients had a Full Scale IQ of 80 or higher (WAIS-IV, The Psychological Corporation), (Wechsler, 2008). Additional details on study entry criteria, including the age, gender matched healthy controls (HC’s, n=24) are provided in Supplement Section 1. The demographic and clinical characteristics of the experimental groups are presented in Table 1.

**Table 1.**
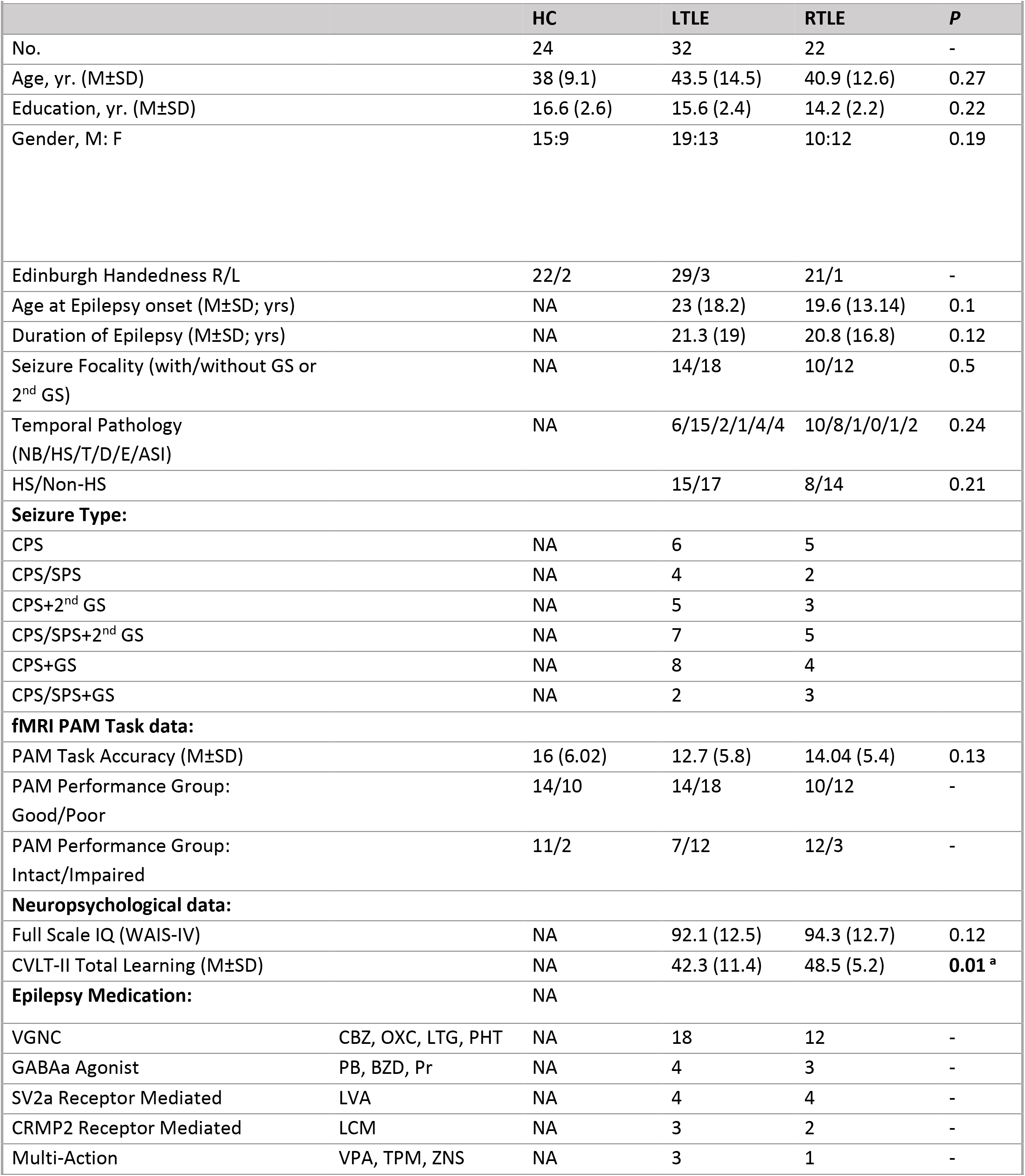
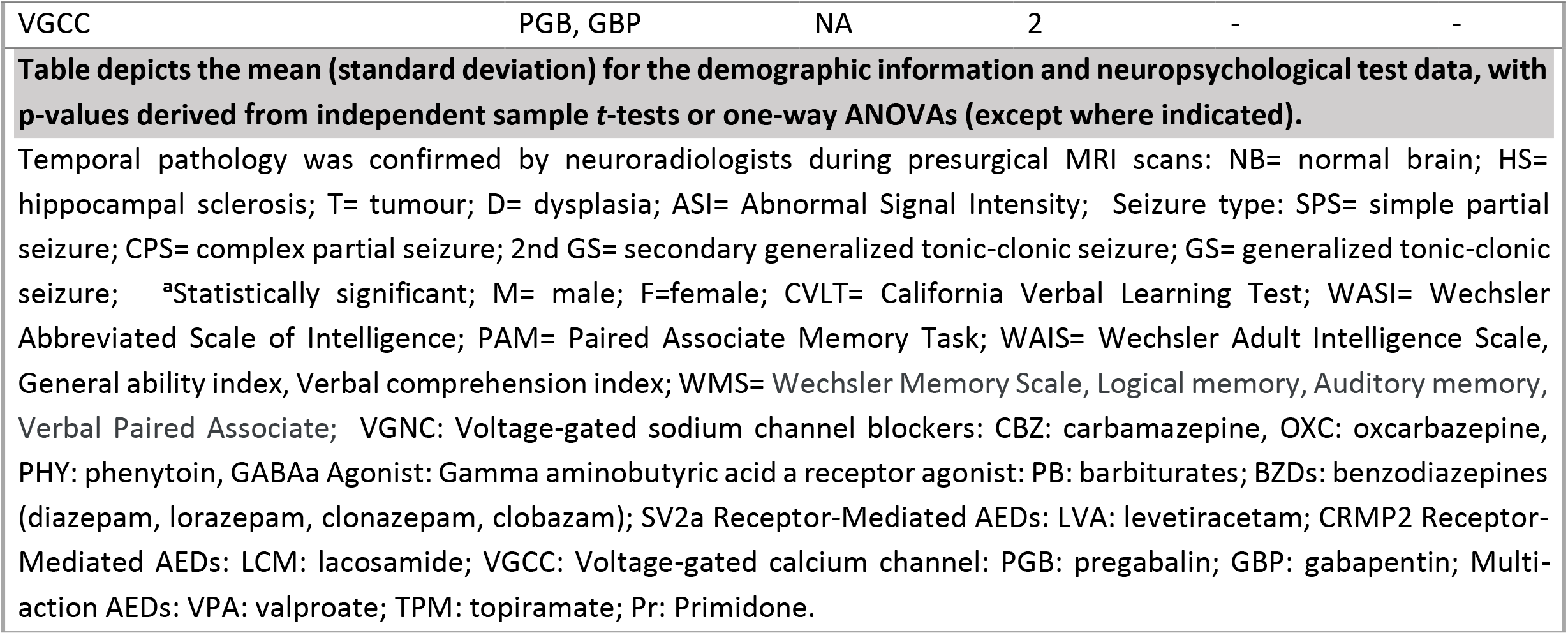
Sample demographic, clinical, PAM task, and neuropsychological data.

### Paired associate memory task

To capture both deleterious memory effects and the potential neuroplastic responses to maintain effective episodic encoding and recall, we chose to trigger memory-relevant BOLD activation through a paired associate memory paradigm (PAM), based upon items and modifications of the Wechsler Memory Scale (Wechsler, 2009). The PAM task design comprised three phases: encoding, when the word pairs are learned and memorized; distraction, when a set of intervening arithmetic problems are solved; cued recall, when one member of the target pair is presented and the participant must speak aloud the second member of the pair. The task design is described in Supplement Section 2, Figures 1 and 2.

**Figure 1.**
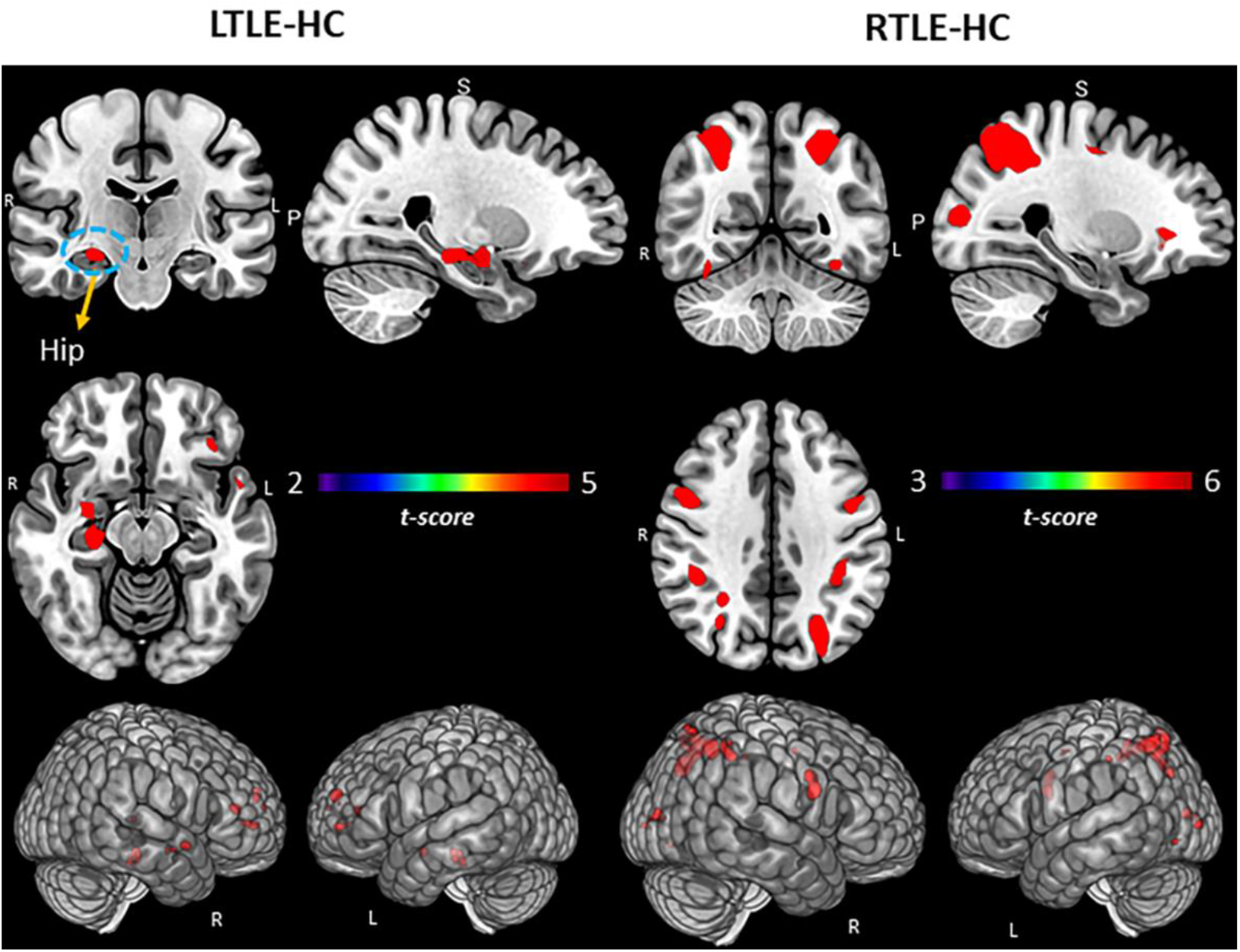
Group comparisons for TLE and HCs. Regional activation associated with subsequent memory effect (SME) minus math condition contrast. Surface rendering and slices in vertical panels show significant activation for LTLE versus HC and RTLE versus HC groups. *pFDR* corrected t-statistic <0.05, cluster level. Hip: Hippocampus.

**Figure 2.**
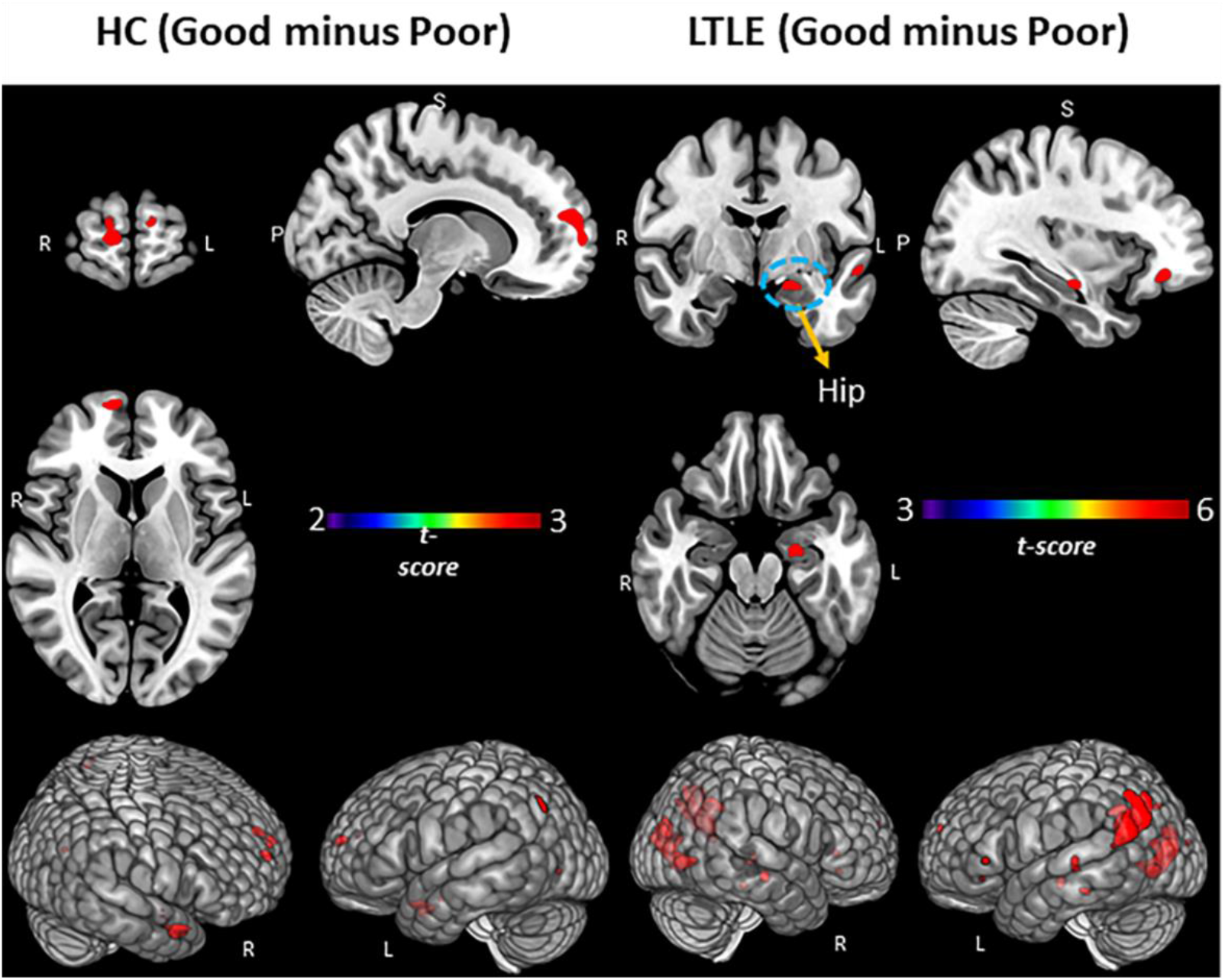
PAM performance (Good versus Poor) within the LTLE and HCs. Regional activation associated with subsequent memory effect (SME) minus math condition contrast. Surface rendering and slices in vertical panels show activation for Good minus Poor subgroups of HC and LTLE. *pFDR* corrected t-statistic <0.05, cluster level. Hip: Hippocampus.

The word pairs that were successfully recovered during the cued recall period were considered successfully encoded. The cued recall responses of forgotten and successfully encoded pairs for each subject were used to calculate the subsequent forgetting (SFE) and subsequent memory (SME) effects. SME’s involved PAM encoding trials subsequently remembered, with SME trials analyzed only for those participants with 8 or more trials remembered (at least 32% recall rate). The activations associated with these SME trials minus the math control condition constituted the main contrast of interest. SFE trials involved encoding trials not subsequently remembered, with the SFE minus math contrast analyzed only for those participants with 8 (32%) or more forgotten trials. Participants with an SME of 60% or better (proportion correctly recalled, at least 16 of 25 pairs correct) were classified as Good PAM performers; participants with lower SME scores were classified as Poor PAM performers. Within the TLE group, we developed a separate grouping strategy to identify performances at the more extremes of the accuracy continuum, considering those with an SME of 70% or better to have displayed fully intact and compensated recall, and those with an SME of 30% or lower to have displayed impaired (uncompensated) recall performance. Note, the HC and RTLE patients generally performed better on the PAM task, with few individuals below 70% in terms of recall accuracy (see Table 1), creating too large an imbalance to make Experimental Group comparisons possible using this intact/compensated versus impaired/uncompensated distinction.

### MRI data acquisition and fMRI preprocessing pipeline

The fMRI scan was obtained on a 3 T Philips Achieva MRI scanner for all participants using an 8-channel head coil. Participants were instructed on all phases of the PAM task and response requirements before entering the scanner. FMRI data preprocessing for PAM task was performed on each subject using fMRIPrep 1.5.8 (Esteban *et al.*, 2019) based on Nipype tool 1.4.1 (Gorgolewski *et al.*, 2011). Details of the MRI acquisition parameters, and fMRIprep pipeline are provided in Supplement Section 3.

### General linear model statistical analyses

Statistical analyses were conducted using (IBM^®^ SPSS^®^ v24), with alpha level set at p <0.05 for multiple comparisons. T-tests or chi-square tests were used, as appropriate, to determine differences in our experimental groups on demographic/clinical characteristics, IQ, and a neuropsychological memory measure available on the TLE patients (Table 1). Utilizing SPM12 (Friston *et al.*, 2011), one or two sample t-tests were carried out to determine whole brain significant activations emerging from the contrast of interest (SME minus math control), either within or between experimental (LTLE, RTLE, HC) or performance groups (Good minus Poor; Intact minus Impaired) *[pFDR* corrected <0.05, cluster level]. We also analyzed the activation associated with the SFE (SFE minus math condition contrast) and tested for group differences in whole brain activation. A two-factor ANOVA analyzing the above SME contrast utilized Experimental Group (e.g., LTLE, HC) and PAM performance (Good versus Poor) as independent variables, with their interaction the effect of interest (*pFDR* corrected <0.05, cluster level).

### Generalized psychophysiological interaction analyses

We performed gPPI analysis to verify the key PAM task-modulated connectivity effects (SME effect versus math control condition) utilizing the CONN toolbox v17.f (McLaren *et al.*, 2012). Whole brain gPPI was applied with the seed based upon the results of the key GLM analyses noted above (see Supplement Section 4 for gPPI details). The first gPPI model was based on the results of the significant interaction effect from the two-factor ANOVA on the SME contrast (Experimental Group [LTLE versus HC] and PAM performance [Good, Poor]). Our second gPPI model was based upon the significant t-test result comparing the Intact and Impaired LTLE groups on the SME activation. These task modulation-dependent gPPI measures allowed us to determine the strength of functional connectivity (FC) to the seed(s) with the whole brain as the search set (*pFDR* corrected < 0.05, two-tailed).

### Support vector machine models and analyses

A linear SVM algorithm (MATLAB, R2018a, with cross-validation through the leave-one-out method) was used for multivariate classification analysis of Intact versus Impaired memory performance in LTLE. We used as inputs the magnitude of the significant regional activations on the PAM task given by the key ANOVA differences emerging from the Intact/Impaired comparison within LTLE, as well as the functional connectivities associated with a key regional SME effect distinguishing Intact/Impaired status (hippocampi bilaterally). This SVM was done solely within the left TLE group as too few patients in the RTLE group displayed impaired/uncompensated performance. To assess the classification success of the SVM model, we computed receiver operating characteristic (ROC) curves, with associated area under the curve data (AUC, p<0.05). The goal of the SVM model was to select the features from our GLM task activation and gPPI functional connectivity data that best discriminated successful verbal memory.

## Results

### Clinical, demographical, and behavioral results

Our sample clinical, demographic, and behavioral data can be seen in Table 1. Our three experimental groups not differ in age, gender, or educational background. Left handedness was identified in a small number of participants as indexed by the Edinburgh Handedness scale (Oldfield, 1971). Age at epilepsy onset, duration of epilepsy illness, and the rate of mesial temporal sclerosis did not differ in the left and right TLE groups (Table 1). As expected, there was significantly worse performance in the left compared to right TLE group on a neuropsychological measure of verbal learning and encoding, the CVLT-II Total Learning (t[df=52] = −2.4, p<.01).

### Experimental group differences in PAM task activation

We examined the pattern of brain activation associated with the SME contrast within each of our Experimental Groups (Supplementary Section, Figure 3). In HC’s, the significant clusters of activation were predominantly in the dominant left hemisphere involving the left inferior frontal, left hippocampal/parahippocampal gyri, left parietal/occipital, and left lingual regions The RTLE group also showed a predominantly left hemisphere pattern, with left hippocampal, left inferior frontal, and left inferior parietal/occipital activations. In the LTLE group, there was bilateral activation in the hippocampus, middle temporal, inferior frontal, and inferior parietal/occipital regions. The LTLE group showed clear and strong evidence of right hemisphere involvement (middle temporal and inferior frontal), with such activity absent in the other groups. All the above activated regions were statistically significant.

**Figure 3.**
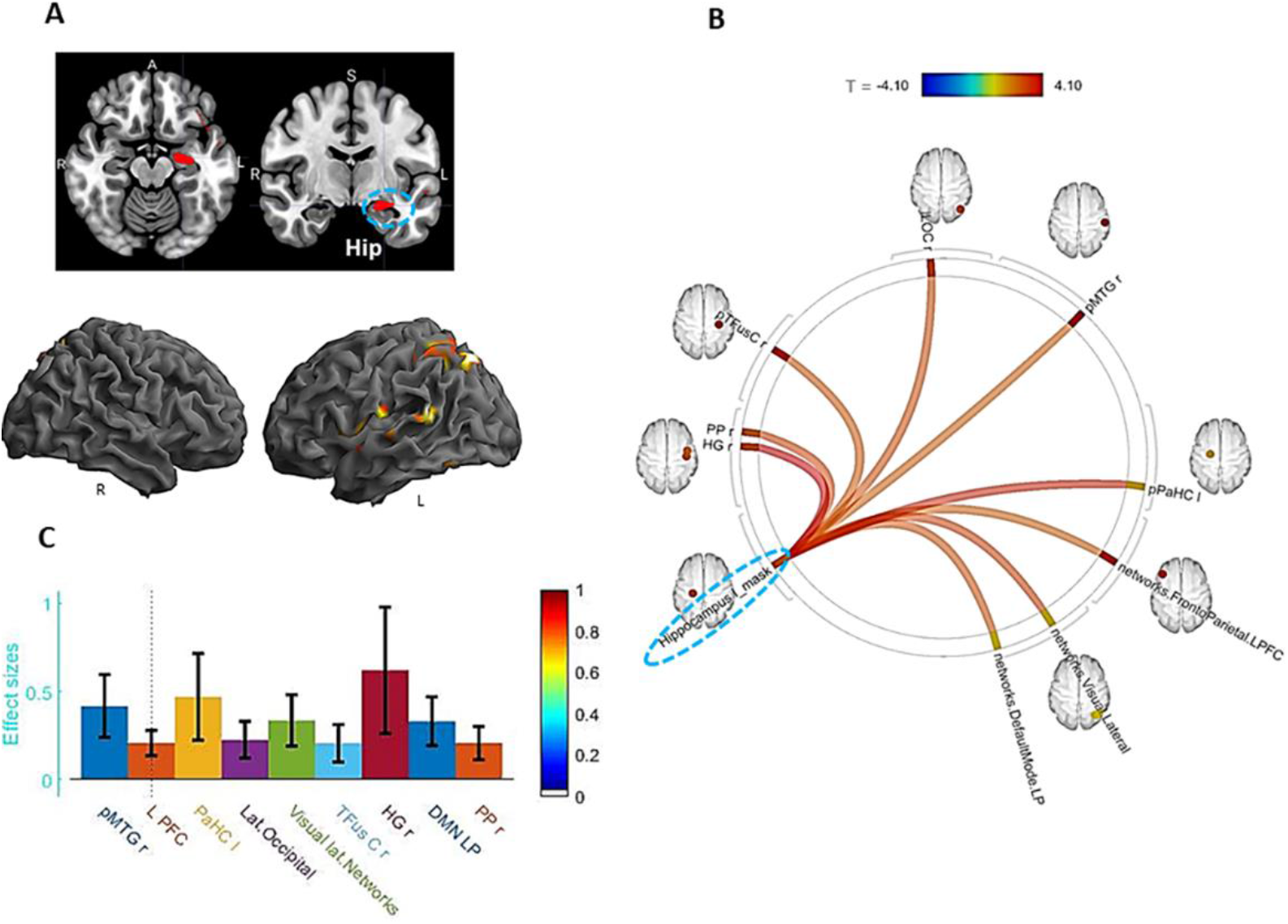
Interaction of experimental (LTLE, HCs) and performance (Good, Poor) groups. **Panel A:** Whole brain surface rendering and slices revealing significant activation resulting from interaction in two-way ANOVA on SME minus Math contrast. Factor One, Experimental Group (LTLE, HCs); Factor 2, PAM performance (Good, Poor; t-statistic *pFDR* corrected < 0.05, cluster level. Results highlight the areas where the SME/Math difference is significantly larger in LTLE patients compared to HCs on the PAM task. **Panel B:** Functional connectivity results of gPPI using left hippocampal region emerging from the GLM interaction effect (see panel A) as seed. Seed cluster (MNI coordinates of maxima: - 28 −24 −16; 103 voxels), involving the time course of the SME minus Math contrast was extracted and the generated gPPI regressor effects (gPPI_β) are displayed. Results revealed nine target regions with functional relationship of statistically significant strength with left hippocampal seed (*pFDR* t-statistic corrected < 0.05). **Panel C:** Bar diagram depicts effect size (mean beta weights from gPPI) for each functional relationship between the source seed (left hippocampus) and target region.

In terms of experimental group differences involving the SME (see Figure 1; Supplementary Section Table 1), compared to HC’s, the LTLE group showed increased activation in the right hippocampus/parahippocampus, along with increased activity in the bilateral middle/superior frontal region. In contrast, compared to HC’s, the RTLE group showed a limited set of differences, with the areas of increased activation involving bilateral posterior/superior regions, parietal cortex, and the occipital lobe.

We also examined whether the experimental groups differed with regard to the SFE minus math contrast. Neither the LTLE nor the RTLE groups differed from HC’s in this regard, suggesting that in contrast to the group differences that emerged for successful memory encoding, the functional neuroanatomy associated with forgetting is similar in our TLE groups relative to controls.

### PAM task performance group activation differences by GLM

We examined the unique patterns of SME-related activations associated with Good versus Poor performance within each of our Experimental Groups (see Table 2A and B, Figure 2). In the HC’s, compared to the Poor performers, the Good PAM Performers showed activations in bilateral anterior/superior frontal, left superior parietal, as well as right anterior/middle temporal lobe. In LTLE, compared to the Poor PAM performers, the Good performers showed several left sided activation increases (hippocampal, supramarginal, and superior temporal, superior parietal. Within RTLE, no reliable activation differences were present between the Good and Poor performance groups.

**Table 2.** MNI coordinates of significant cluster maxima for subsequent memory effect minus math contrast demonstrating performance group differences within HC and LTLE.

| Region | <u>Peak MNI Coordinates</u> |  |  |  |
| --- | --- | --- | --- | --- |
|  | k | x | y | z |
| **A. HC (Good minus Poor)** |  |  |  |  |
| R Anterior/superior frontal gyrus | 108 | 14 | 60 | 16 |
| L Anterior/superior frontal gyrus | 78 | -6 | 56 | 16 |
| R Anterior/middle temporal gyrus | 83 | 40 | 4 | -32 |
| L Superior parietal gyrus | 34 | -24 | -68 | 58 |
| R Frontal pole | 26 | 22 | 58 | -2 |
| **B. LTLE (Good minus Poor)** |  |  |  |  |
| L Supramarginal gyrus | 248 | -62 | -52 | 20 |
| L Superior temporal gyrus | 177 | -44 | -24 | 6 |
| L Precuneus | 81 | -16 | -56 | 34 |
| L Superior parietal lobule | 60 | -32 | -54 | 38 |
| R Angular Gyrus | 46 | 38 | -56 | 28 |
| L Hippocampus | 43 | -36 | -22 | -8 |
| Subpeaks of the interest are also included. The cluster thresholds correspond to corrected False Discovery Rate with significance level of height $p < 0.05$. | | | | |

We next sought to determine if the regional activation differences associated with Good versus Poor PAM performances varied as a function of TLE or HC status. We utilized a two-factor ANOVA model, with the interaction between these factors the effect of interest. The results showed that the Good versus Poor SME activation differences did vary as a function of LTLE or HC group (see Table 3A and Figure 3, panel A). More specifically, in LTLE compared to HC’s, the Good PAM performers displayed significantly greater activation than the Poor performers in the left hippocampus, and several left-hemisphere regions (superior parietal, fusiform, posterior superior temporal, central operculum, and Heschl’s gyrus). The interaction effect involving PAM performance (Good versus Poor) and the RTLE and HC groups produced no statistically reliable results.

**Table 3.** MNI coordinates of significant cluster maxima for subsequent memory effect minus math contrast from ANOVA interaction effect and performance group t-test within LTLE.

|  |  |  |  |  |
| --- | --- | --- | --- | --- |
| **Region** | **<u>Peak MNI Coordinates</u>** |  |  |  |
|  | **k** | **x** | **y** | **z** |
| **A. Two-Way ANOVA Interaction, Factor 1 (LTLE, HC) and Factor 2 (Good, Poor)** |  |  |  |  |
| L Superior parietal lobule | 1016 | -16 | -52 | 34 |
| L Fusiform gyrus | 523 | -38 | -74 | -8 |
| L Posterior/superior temporal gyrus | 358 | -46 | -26 | 6 |
| R Thalamus | 322 | 10 | -34 | 8 |
| L Central operculum | 292 | -56 | -4 | 4 |
| L Heschl's Gyrus | 226 | -46 | -26 | 6 |
| L Hippocampus | 103 | -28 | -24 | -16 |
| **B. T-test, LTLE PAM Groups (Intact minus impaired)** |  |  |  |  |
| R Posterior superior temporal gyrus | 609 | 58 | -58 | 14 |
| R-Heschl's Gyrus | 377 | 38 | -28 | 6 |
| L Supramarginal/angular gyrus | 215 | -40 | -70 | 40 |
| R Cerebellum | 177 | 16 | -62 | 10 |
| L Hippocampus | 75 | -20 | -24 | -18 |
| R Hippocampus | 72 | 14 | -22 | 14 |
| Activated cluster description for Interaction; A) LTLE and HC groups, B) LTLE (Intact versus Impaired) group. Subpeaks of the interest are also included. The cluster thresholds correspond to corrected False Discovery Rate with significance level of height threshold ($p < 0.05$). | | | | |

To delineate cognitive compensation in our TLE patients, we examined the SME effects associated with the more extreme ends of PAM performance within our TLE groups. These results showed that in LTLE, clearly Intact compared to Impaired performance was associated with increased activation in both the left and the right hemispheres (see Table 3B; also Figure 4, panel A). Importantly, we observed bilateral hippocampal activation, but also right hemisphere activation (posterior superior temporal, Heschl’s gyrus, cerebellum), and left angular gyrus.

**Figure 4.**
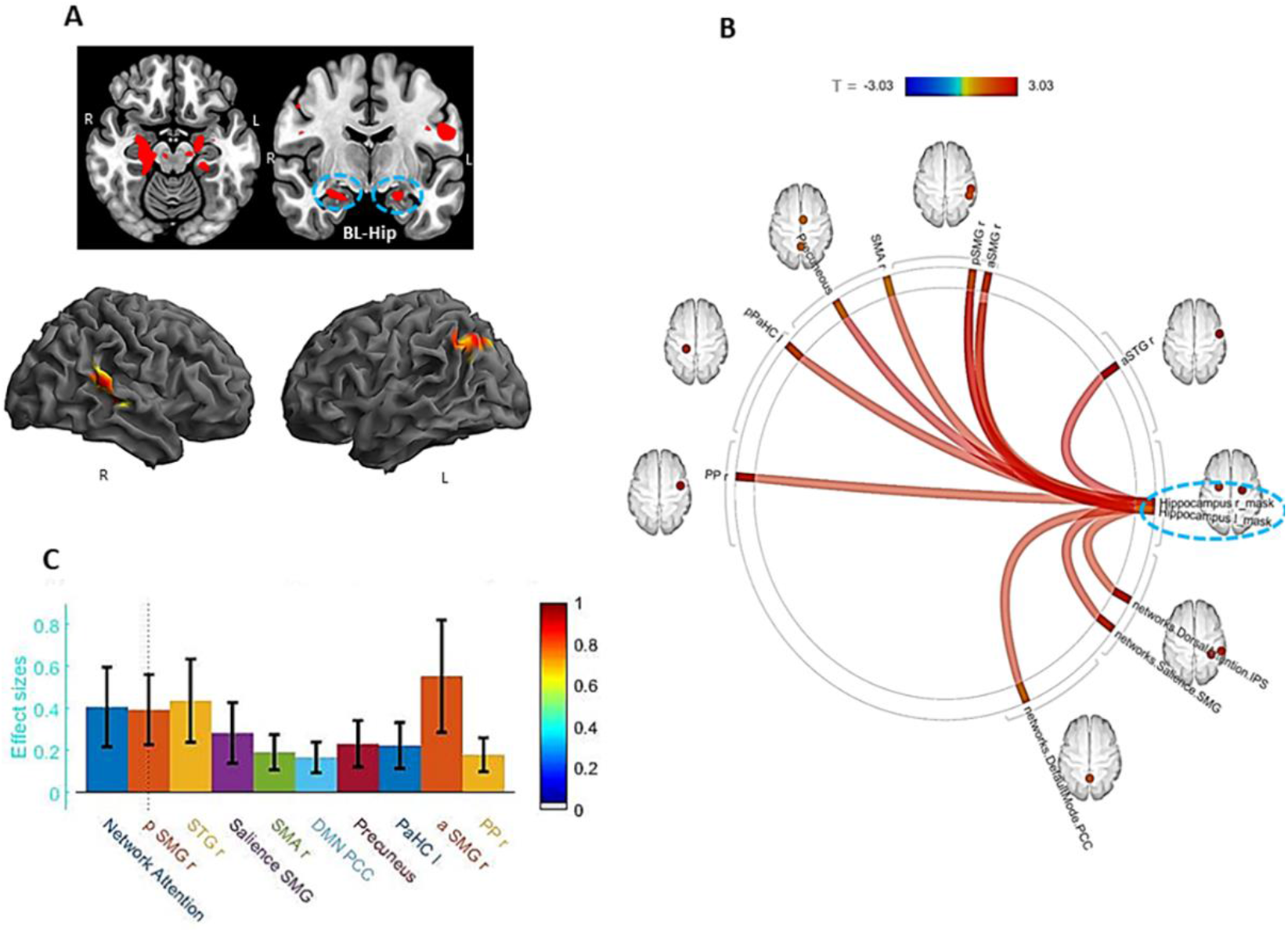
Sub-group comparison (Intact minus Impaired) within LTLE. **Panel A:** Whole brain surface rendering and slices revealing significant activation (SME minus Math contrast) resulting from t-test of sub-group difference (intact minus impaired) within the LTLE patients (*pFDR* t-statistic corrected, <0.05, cluster level). Results highlight the areas where the SME/Math difference is significantly larger in the LTLE patients with intact compared to impaired performance levels on the PAM task. **Panel B:** Functional connectivity results of gPPI using bilateral hippocampal region emerging from the t-test effect (see panel A) as seed. Seed clusters of left (MNI coordinates of maxima: −20 −24 −18; 75 voxels) and right (14 −22 14; 72 voxels) hippocampus, involving the time course of the SME minus Math contrast was extracted and the generated gPPI regressor effects (gPPI_β) are displayed. Results revealed 10 target regions with functional relationship of statistically significant strength with combined hippocampal seed (*pFDR* t-statistic corrected < 0.05). **Panel C:** Bar diagram presented effect size (mean beta weights from gPPI) for each functional relationship between the source seed (combined left and right hippocampus) and target regions.

### Functional connectivity effects (gPPI) with key GLM results as seed

Having established that activation in the left hippocampus is a key region distinguishing Good and Poor performances in LTLE compared to controls (see Table 3A), and that activation in the bilateral hippocampus is important to the Intact and Impaired distinction within LTLE (Table 3B), we next sought to determine if there are functional connectivities to these key areas that serve as mediators of successful memory during the PAM task (i.e., mediate SME trials but not the math control condition). To accomplish this, we utilized gPPI, focusing on the whole brain connectivity related to the hippocampal seed given by the above GLM analyses.

With regard to the Good versus Poor difference in LTLE and HC, we found strong functional connectivity between 9 regions and the left hippocampal seed during the SME trials, as presented in Figure 3, Panels B and C. These regional FC’s involved mostly right-sided regions (planum polar, Heschl’s gyrus, temporal fusiform, lateral occipital, posterior middle temporal gyrus). Other regions that showed increased connectivity to the left hippocampus were the left parahippocampus and parts of well-established intrinsic FC networks such as the default mode (right lateral parietal), visual lateral (right side), and fronto-parietal (left prefrontal cortex) networks. Of these regions, the right Heschl’s, right posterior middle temporal, and left parahippocampal regions showed the strongest FC with the left hippocampus.

With regard to the Intact versus Impaired difference within LTLE, we found 10 regions that showed strong functional connectivity to the bilateral hippocampi during the SME trials (see Figure 4, Panels B, and C). These regional FC’s involved mostly right hemisphere regions (planum polar, anterior/posterior supramarginal, superior temporal, and supplementary motor area), in addition to the left parahippocampus. Other regions connected to the bilateral hippocampi were members of well-established intrinsic FC networks such as the dorsal attention (right inferior parietal sulcus), salience (right supramarginal gyrus), and default mode (precuneus, posterior cingulate cortex) networks. The right anterior/posterior supramarginal gyrus, right superior temporal gyrus, and dorsal attention network region (right inferior parietal sulcus) showed the strongest FC with the bilateral hippocampi.

### SVM prediction of Intact/Compensated versus Impaired/Uncompensated status in LTLE

To further understand the features in our data that best discriminated intact from impaired performance in LTLE, we utilized SVM multivariate classification analyses focused on determining whether SME-related activation magnitudes (see Table 4, effect size variables) or the strength of FC’s with the bilateral hippocampi seed used in our gPPI model (see Table 4, FC coefficient variables) were better classifiers. For the PAM task, six variables produced statistically significant (p<.05) AUC values of .796 or higher, mostly involving right hemisphere variables. Four of these six variables involved SME activation effects (right hemisphere -- superior temporal, Heschl’s gyrus; left hemisphere – hippocampus, angular gyrus), with two FC variables reaching statistical significance as a classifier (right anterior supramarginal and right planum polar gyri).

**Table 4.** Multivariate classification analysis (SVM) within LTLE utilizing significant activation effect sizes and gPPI functional connectivity coefficients (with Seed) as input.

Model: Response Variable: Intact minus Impaired Status Based on PAM Task
| <u>Region (Effect Sizes)</u> | <u>CA (%)</u> | <u>AUC</u> | <u>SE</u> | <u>P-Value</u> | <u>95% CI</u> |
| --- | --- | --- | --- | --- | --- |
| RHCP | 54 | .633 | .154 | .406 | .330-.935 |
| RSTG | 78.2 | .898 | .085 | .013* | .732-1.0 |
| LHCP | 76.1 | .878 | .096 | .018* | .689-1.0 |
| LAG | 81.2 | .959 | .050 | .004* | .861-1.0 |
| RHG | 79 | .939 | .067 | .006* | .807-1.0 |
| RCBLM | 56 | .512 | .119 | .910 | .278-.746 |
| <u>Region (FC Coef.)</u> |  |  |  |  |  |
| aSTGr | 54.2 | .551 | .167 | .749 | .224-.878 |
| aSMGr | 74.3 | .796 | .123 | .045* | .554-1.0 |
| pSMGr | 53 | .510 | .164 | .949 | .188-.832 |
| SMAr | 52.2 | .510 | .164 | .949 | .188-.832 |
| Precuneus | 51 | .490 | .167 | .949 | .163-.816 |
| pPaHCl | 49.8 | .408 | .168 | .565 | .078-.738 |
| PPr | 78.2 | .918 | .075 | .009* | .772-1.0 |
| DMN.PCC | 46.2 | .388 | .159 | .482 | .077-.699 |
| Networks.Salience.SMG | 46 | .327 | .155 | .277 | .022-.631 |
| Networks.Attention.IPS | 54 | .531 | .168 | .848 | .201-.860 |
| *Statistically significant; CA: Classification accuracy; AUC: area under curve; SE: standard error; CI: confidence interval. |  |  |  |  |  |

Lastly, we examined whether the above subset of significant variables remained reliable and effective classifiers after including FSIQ or neuropsychological memory (CVLT-II Total Learning) in the model and re-running the identical SVM. The results showed that the above six variables remained significant in the SVM. FSIQ was not a significant predictor (AUC, p-value=.48), however, the neuropsychological measure did emerge as significant (AUC, p-value<.05), suggesting that our Intact/Impaired grouping was consistent with the baseline memory levels of our LTLE patients. We examined in LTLE the correlation between our SVM predictor set (16 variables) and age at epilepsy onset and duration of epilepsy. Duration bore no significant relations, but age of epilepsy onset was negatively correlated with two variables, the left precuneus (r=-.56, p<.05) and right planum polar area (r=-.59, p<.05). These findings indicated that earlier epilepsy onset was associated with stronger FC modulation between those two regions and the bilateral hippocampi seed. Age of onset was not significantly different between the Intact/Impaired groups (Intact, onset=14.7 years; Impaired, onset=23.1 years). Nonetheless we ran the identical SVM and found the same six variables remained significant in terms of classifying our Intact/Impaired LTLE patients, with age of onset also emerging as a significant classifier (AUC=.84, p<.05).

## Discussion

We utilized both activation magnitude and functional connectivity to demonstrate the regional mechanisms involved in the reorganization of episodic memory in temporal lobe compromised epilepsy patients. We carefully linked our fMRI analyses to successfully encoded words through the use of SME effects. SME contrasts reflecting activation relative to a math control condition did show reliable regional effects within each of our experimental groups, but an investigation of unsuccessful PAM performance (SFE) showed no reliable effects in any of our groups (HC, LTLE, or RTLE). The latter makes clear that merging these distinct cognitive performances (trials) likely hides the unique computational properties related to compensatory, successful memory reorganization. Our data showed that left, but not right, TLE differed from healthy controls in their brain maps of PAM SME’s, with the maps of the LTLE patients more distinctly right-sided, particularly in mesial temporal areas, as well as showing bilateral superior frontal activation compared to controls.

The above findings, however, did not take into account performance levels. Indeed, most unique to our study is the examination of different memory performance levels, captured by categorizing our participants as Good versus Poor PAM performers, and, in the case of our TLE patients, as clearly Intact versus Impaired. Several distinctive features of Good performance in LTLE relative to HC were present. For instance, there was increased activation in left mesial (left hippocampal) and several left posterior regions (superior parietal and temporal, fusiform, central operculum, and Heschl’s gyrus; see Table 3A, Figure 3a). These left hemisphere areas seemed to be uniquely and abnormally upregulated among the Good LTLE performers. We found no reliable Good versus Poor activation differences within RTLE, or when comparing RTLE to HC’s.

An added layer of our investigation of performance effects involved use of a stringent criterion, one that separated performance levels so as to be sure to isolate truly intact versus impaired recall levels relative to same age peers (SME at 70% accuracy or better; 30% or worse). This distinction showed that, indeed, a different set of regions were responsible for these higher, truly compensated performance levels. Intact as compared to Impaired performance was associated with bilateral hippocampi, left supramarginal/angular gyrus, right posterior superior temporal gyrus, and other right-sided areas (Heschl’s, cerebellar) of activation. These data made clear that in contrast to Good performance levels, Intact or heightened performance levels were strongly right hemisphere in nature, involving uniquely increased activity in the (language) non-dominant hemisphere contralateral to the ictal focus.

These more stringent performance-based changes were unique to LTLE. For instance, while the RTLE group did show some PAM activation differences relative to HC’s in bilateral parietal areas (see Figure 1; Supplement Section Table 1), both RTLE and HC’s showed a similar strong left hemisphere pattern, involving the left hippocampus and left inferior frontal regions (see Supplement Section Figure 3). On the PAM task, the HC’s brain regional involvement did also vary with performance. Good, advantaged memory performance was characterized by bilateral activity involving the anterior/superior frontal lobe, left superior parietal, and right anterior/ middle temporal regions (Table 2, Figure 2). A conjunction analysis of Good performers in the HC and LTLE groups showed a common left-sided activation pattern (hippocampus, inferior frontal, middle temporal, superior temporal, and left angular gyrus, lingual gyrus), indicating that some of the ipsilateral effects observed in LTLE are, indeed, normative and not unusual brain responses. An area notable because of its very limited presence in the normative responses of the Good and Intact LTLE groups involved frontal cortex, as the frontal lobe was a prominent part of the Good performance response in HC’s.

Our data also made clear that most of these shared (i.e., normative) areas of activation differed from and, therefore, were not critical to Intact/Compensated, or even Good performance in LTLE. In brief, the advantaged areas in the HC’s were more anterior/middle temporal and frontal in location compared to the advantaged LTLE performers. With the above in mind, we can see that in terms of activation magnitude the areas of increased activity in the clearly intact/compensated LTLE patients that are non-normative involved the right hippocampus response, the right posterior superior temporal gyrus, and the other right hemisphere clusters, responses clearly missing in the other groups. Note, the less accurate, but Good, memory performances in LTLE that were non-normative involved left superior parietal, left fusiform, left central operculum, left Heschl’s gyrus, and the right thalamus. With the exception of the right hippocampus, which has been observed in normals during paired associate paradigms (Clark *et al.*, 2018; Gould *et al.*, 2003), the non-normative areas we found to be unique to the LTLE Good and Intact performers are not considered primary computational regions for episodic memory (i.e., implementing associative encoding, memory consolidation). This unique profile of non-normative areas suggested these represent forms of computational support, as opposed to areas of computational primacy, for the production of successful episodic memory. Accordingly, our data provided strong evidence that when seeking to clarify the regions most unique to implementing successful, truly compensated memory reorganization in LTLE, it is important to distinguish between performance levels.

To more specifically characterize the brain features associated with intact/compensated status in LTLE, we looked not just at PAM activation magnitude, but also changes in functional connectivity unique to successful PAM encoding. The goal of these gPPI analyses was to find the areas communicating during encoding with the key regions present in our activation magnitude data, taking advantage of our knowledge that such regions (hippocampi) play a critical role in primary memory computations based upon a literature dating back decades (Lech and Suchan, 2013; Scoville and Milner, 1957; Squire and Zola-Morgan, 1991). Any observed significant FC’s can be said to reflect regions with activity synchronized and correlated with the hippocampus seed, implying that they mediate successful memory, even though their activation magnitude may not have been sufficient to appear in our GLM findings. The results revealed that nine regions had reliable FC with the left hippocampus in Good relative to the Poor group in LTLE compared to HC’s (see Figure 3a). These regional FC’s implicate extensive SME-related communication between the left hippocampi and right-sided regions, in addition to contact with regions that are part of intrinsic networks known either to be associated with memory processes (default mode), or other processes often associated with a strong memory response to viewed material (executive function, fronto-parietal network; visual lateral network). These results stand in contrast to the activation magnitude data which showed that the Good/Poor groups in the LTLE and HC’s differed predominantly in left-sided areas.

In our attempt to understand the communication circuitry underlying truly intact/compensated status in LTLE, our gPPI model (bilateral hippocampal seed) revealed 10 regions with reliable FC during encoding. Consistent with the right-hemisphere nature of the PAM activation magnitude data, we saw mostly right-sided FC’s associated with Intact status. Perhaps not unexpectedly, the bilateral hippocampi showed connectivity to regions that are part of intrinsic functional networks well-associated with either memory encoding processes (e.g., the default mode network, precuneus, posterior cingulate; see our work (Doucet *et al.*, 2014), or other cognitive computations that can be seen as important for a successful memory (e.g., dorsal attention and salience networks). These FC’s gave us a window through which to view the broader networks supporting compensated memory in these temporal lobe compromised patients. As a test of whether these FC’s were in any sense normative, we conducted the identical gPPI model in HC’s (bilateral hippocampal seed). The resulting gPPI was not significant (*pFDR* corrected, <.20), with very different mediating regions in the model. This follow-up analysis demonstrated that the 10 regions mediating memory performance in the Intact group did not reflect normal increases in FC during the SME trials.

Our performance focused gPPI’s were similar in that both implied strong communication with the non-ictal, non-dominant right temporal areas in the advantaged LTLE groups, as well as strong connections to the same intrinsic connectivity network, the DMN, known for its association with episodic memory. Both gPPI’s also showed connectivity to the left, ipsilateral parahippocampus. The two gPPI’s, however, differed with respect to the other intrinsic networks recruited to support successful memory (the fronto-parietal network for distinguishing Good versus Poor LTLE status; the dorsal attention and salience networks for the separation of Intact/Impaired status). The literature does suggest that these three intrinsic networks bear some functional similarity. For instance, each of these networks involve top-down responses that are phasic forms of cognitive control, each is triggered by exogenous stimuli, and each evokes a cognitive goal that establishes what stimuli are important, with subsequent initiation and maintenance of task focus and engagement (Sadaghiani and D’Esposito, 2015). This complex, but shared functionality, can be viewed as set of facilitative communications that support episodic memory. It is important to be reminded that many of the regions forming these memory-related connectivities with the hippocampus were not activation hotspots in the linked PAM-activation data.

Lastly, having established that distinct performance levels produced different task-related activations and memory-mediated functional connectivities, we sought to determine which feature(s) best distinguished Intact versus Impaired status in LTLE, noting that this grouping provided the most compelling contrast to argue that our findings truly represented compensatory memory reorganization. Utilizing AUC values from the SVM model as our guide (see Table 4), we found that activation magnitudes were generally better classifiers of Intact/Impaired status on the PAM, with these regions involving a mix of right and left hemisphere structures. The reliable FC predictors involved right hemisphere-based connections (right anterior supramarginal gyrus, right temporal planum polar) to the bilateral hippocampal seeds. Looking across these predictors, it is really the pattern involving right posterior temporal (magnitudes: superior temporal gyrus, Heschl’s; FC’s: temporal planum polar) and nearby right inferior parietal features (FC: supramarginal gyrus) that was most striking. Thus, our SVM results clearly demonstrated the importance of right hemisphere structures in compensatory memory responses in LTLE, making clear that with the exception of the left hippocampus, the critical predictors of Intact/Compensated status were not homologues of the left hemisphere areas known to associated with primary episodic memory computations in normals.

A potential implication of our results is that truly compensatory memory reorganization in LTLE can be understood to take two forms. One form involves change to the regions conducting *primary memory computations* such as associative learning, memory engram formation and consolidation. Relevant to this, our SVM data indicated that no fundamental reorganization occurred with regard to *computational primacies*, as left hippocampal SME-related activation remained one of the best predictors of Intact/Compensated status. The second form of reorganization involves change to the regions providing *computational support* to the core and primary memory processes. These support regions reflect recruitment of cognitive functions that facilitate, and perhaps even are a necessity, for strong, highly successful memory performance. Relevant to this, our SVM data indicated that there were non-normative regional involvements mostly in the right temporal lobe providing *computational support* for memory processing either through task-relevant upregulations in activity or task-mediated increases in connectivity to the bilateral hippocampi. One might speculate as to the cognitive nature of these supportive functionalities (e.g., dedication of attentional resources, cognitive control/strategy, engagement of task-relevant stimuli, working memory, increased reliance on lexical association or search functions, auditory processing), but the fact that none of these areas discriminated Good versus Poor performance in HC’s suggested the regions identified by our SVM model were unique to the implementation of successful memory in LTLE. Our PAM activation magnitude and FC data pointed to other regional differences between the Intact/Impaired groups, providing additional clues to the functionalities that may have provided computational support for strong, successful episodic memory in LTLE (see (Tracy and Osipowicz, 2011) for discussion of alterations in cognitive networks that might drive adaptive performance in the face of disease). Ultimately, however, our data remained silent on the specific role played by the regions we have identified as strongly associated with compensated memory organization in LTLE.

Several methodologic considerations are pertinent to mention. Follow up examination revealed that our SVM model is capturing activation magnitude and FC effects that cannot be accounted for by cognitively relevant baseline characteristics of our LTLE sample (IQ, neuropsychological memory), or clinical characteristics such as anti-epileptic medication, epilepsy duration, or age or illness onset. Interestingly, our data does point to the possibility that earlier onset for LTLE lays down a set of increased connectivities not just to the ictal but also the non-ictal hippocampus. Other methodologic and conceptual considerations are noted in Supplement Section 6. These involve the likely importance of white matter changes to cognitive reorganization, and the potential relevance of task-mediated communication changes with other regions, not just the hippocampi. Also, in describing forms of memory reorganization, while we noted several hemispheric patterns, we emphasize that such reorganization processes must be understood at the functional/regional, not hemispheric level. Lastly, we acknowledge that we did not capture individual patterns of reorganization, and the forms of compensatory reorganization we describe for LTLE may not hold for all cognitive processes/tasks, nor for all forms of epilepsy.

Our data showed that the brain reorganizations implementing Intact/Compensated versus Impaired/Uncompensated memory performance in LTLE reflects a complex substrate. The theme throughout is that non-ictal, non-dominant posterior temporal regions are most important, recruiting both increased regional activity (posterior superior temporal, Heschl’s gyrus) and increased modulatory communication with the hippocampi, all features that are missing in the Impaired/Uncompensated LTLE patients. The right hemisphere areas that emerged as most important are neither contralateral homologues to left hemisphere areas, nor are they areas traditionally considered computationally primary for episodic memory. Yet, the story is complex as activation increases in ictal-side areas also appear to be necessary recruitments. Importantly, none of these areas of increased activation, nor FC’s, were associated with advantaged (Good) episodic memory in healthy controls, making clear their unique association with compensatory memory status in LTLE.

Our emphasis on different performance levels yielded insight not just into the regional changes in activation magnitude and FC that are most crucial to compensated episodic memory, but makes clear that in doing so different forms of cognitive reorganization emerge and should be distinguished. Namely, regions can reflect a change in *computational primacy*, and in this respect our Intact/Compensated LTLE group showed little change relative to HC’s as the hippocampi and left parahippocampus, as well as regions that are part of an intrinsic network linked to memory (the default mode), remained important distinguishing features. Our performance effects also revealed a set of regions that likely provided *computational support*, with our data demonstrating that in this respect our Intact/Compensated LTLE did show differences, i.e., adaptive abnormalities, relative to HC’s, involving mostly the right posterior temporal lobe, as well as regions that are members of intrinsic networks that could readily provide functionalities to enhance memory through increased communication with the bilateral hippocampi.

Our data provides evidence of changes in computational support, as opposed to computational primacies, not seen in healthy controls and missing in performance disadvantaged LTLE groups. In so doing, we isolated unique regional activations and mediating FC’s that implement truly compensatory reorganization in LTLE. Our results provide a new perspective of memory deficits in TLE, as it is not just a knockout of a key functional hub such as the hippocampus that causes deficits. Deficits may also arise from a failure to instantiate a complex set of reorganization responses capable of preserving memory. Such responses, whether they involve increases in regional activation or increases in FC with core memory structures, provide the computational support to ensure effective memory performance. Most important, our results increase our understanding of adaptive brain responses and plasticity in epilepsy.

## Abbreviations

PAM: Paired Associate Memory Task
FC: Functional Connectivity
gPPI: Generalized Psychophysiological Interaction
SVM: Support Vector Machine
HC: Healthy Control
LTLE: Left Temporal Lobe Epilepsy
RTLE: Right Temporal Lobe Epilepsy
SME: Subsequent Memory Effect
SFE: Subsequent Forgetting Effect

## Acknowledgements

The authors thank all the healthy controls and patients with epilepsy, kept anonymous, who provided their time to participate in this study. J.I.T. acknowledges funding support from the National Institute of Health, National Institute of Neurological Disorders and Stroke, R01 NS112816-01. M.R.S. acknowledges funding support from the National Institute of Health, R01 NS106611 and U01 NS113198.

## Notes

**Disclosure:** None of the authors have any conflict of interest to disclose.

### Competing Interest Statement

The authors have declared no competing interest.

## References

Alessio A, Pereira FR, Sercheli MS, Rondina JM, Ozelo HB, Bilevicius E, et al. Brain plasticity for verbal and visual memories in patients with mesial temporal lobe epilepsy and hippocampal sclerosis: an fMRI study. Hum Brain Mapp 2013; 34: 186–99.

Baxendale S, Thompson P, Harkness W, Duncan J. Predicting memory decline following epilepsy surgery: a multivariate approach. Epilepsia 2006; 47: 1887–94.

Bell BD, Davies KG. Anterior temporal lobectomy, hippocampal sclerosis, and memory: recent neuropsychological findings, Neuropsychol Rev 1998; 8: 25–41.

Bonelli SB, Powell RH, Yogarajah M, Samson RS, Symms MR, Thompson PJ, et al. Imaging memory in temporal lobe epilepsy: predicting the effects of temporal lobe resection. Brain 2010; 133: 1186–99.

Bonelli SB, Thompson PJ, Yogarajah M, Powell RH, Samson RS, McEvoy AW, et al. Memory reorganization following anterior temporal lobe resection: a longitudinal functional MRI study. Brain 2013; 136: 1889–900.

Bowles B, Crupi C, Pigott S, Parrent A, Wiebe S, Janzen L, et al. Double dissociation of selective recollection and familiarity impairments following two different surgical treatments for temporal-lobe epilepsy. Neuropsychologia 2010; 48: 2640–7.

Burke JF, Long NM, Zaghloul KA, Sharan AD, Sperling MR, Kahana MJ. Human intracranial high-frequency activity maps episodic memory formation in space and time. Neuroimage 2014a; 85: 834–43.

Burke JF, Ramayya AG, Kahana MJ. Human intracranial high-frequency activity during memory processing: neural oscillations or stochastic volatility? Curr Opin Neurobiol 2015; 31: 104–10.

Burke JF, Sharan AD, Sperling MR, Ramayya AG, Evans JJ, Healey MK, et al. Theta and high-frequency activity mark spontaneous recall of episodic memories. J Neurosci 2014b; 34:11355–65.

Cheung MC, Chan AS, Lam JM, Chan YL. Pre- and postoperative fMRI and clinical memory performance in temporal lobe epilepsy. J Neurol Neurosurg Psychiatry. 2009; 80: 1099–106.

Clark IA, Kim M, Maguire EA. Verbal Paired Associates and the Hippocampus: The Role of Scenes. J Cogn Neurosci. 2018; 12: 1821–1845.

Coras R, Pauli E, Li J, Schwarz M, Rossler K, Buchfelder M, et al. Differential influence of hippocampal subfields to memory formation: insights from patients with temporal lobe epilepsy. Brain. 2014; 137: 1945–57.

Doucet GE, Skidmore C, Evans J, Sharan A, Sperling MR, Pustina D, et al. Temporal lobe epilepsy and surgery selectively alter the dorsal, not the ventral, default-mode network. Front Neurol. 2014; 5: 23.

Esteban O, Markiewicz CJ, Blair RW, Moodie CA, Isik AI, Erramuzpe A, et al. fMRIPrep: a robust preprocessing pipeline for functional MRI. Nat Methods. 2019; 16: 111–16.

Flugel D, Cercignani M, Symms MR, O’Toole A, Thompson PJ, Koepp MJ, et al. Diffusion tensor imaging findings and their correlation with neuropsychological deficits in patients with temporal lobe epilepsy and interictal psychosis. Epilepsia. 2006; 47: 941–4.

Friston KJ, Ashburner JT, Kiebel SJ, Nichols TE, Penny WD. Statistical Parametric Mapping: The Analysis of Functional Brain Images: The Analysis of Functional Brain Images. Academic Press; 2011.

Gleissner U, Helmstaedter C, Schramm J, Elger CE. Memory outcome after selective amygdalohippocampectomy: a study in 140 patients with temporal lobe epilepsy. Epilepsia. 2002; 43: 87–95.

Glikmann-Johnston Y, Saling MM, Chen J, Cooper KA, Beare RJ, Reutens DC. Structural and functional correlates of unilateral mesial temporal lobe spatial memory impairment. Brain. 2008; 131: 3006–18.

Gorgolewski KC, Burns D, Madison C, Clark D, Halchenko YO, Waskom ML, et al. Nipype: a flexible, lightweight and extensible neuroimaging data processing framework in python. Front Neuroinform. 2011; 5: 13.

Gould RL, Brown RG, Owen AM, ffytche DH, Howard RJ. FMRI BOLD response to increasing task difficulty during successful paired associates learning. Neuroimage. 2003; 20: 1006–19.

Helmstaedter C, Elger CE. Cognitive consequences of two-thirds anterior temporal lobectomy on verbal memory in 144 patients: a three-month follow-up study. Epilepsia. 1996; 37: 171–80.

Khalil AF, Iwasaki M, Nishio Y, Jin K, Nakasato N, Tominaga T. 2016. Verbal Dominant Memory Impairment and Low Risk for Post-operative Memory Worsening in Both Left and Right Temporal Lobe Epilepsy Associated with Hippocampal Sclerosis. Neurol Med Chir (Tokyo), 56: 716–23.

Kim H, Yi S, Son EI, Kim J. Material-specific memory in temporal lobe epilepsy: effects of seizure laterality and language dominance. Neuropsychology. 2003; 17: 59–68.

Law JR, Flanery MA, Wirth S, Yanike M, Smith AC, Frank LM, et al. Functional magnetic resonance imaging activity during the gradual acquisition and expression of paired-associate memory. J Neurosci. 2005; 25: 5720–9.

Lech RK, Suchan B. The medial temporal lobe: memory and beyond. Behav Brain Res. 2013; 254: 45–9.

Limotai C, McLachlan RS, Hayman-Abello S, Hayman-Abello B, Brown S, Bihari F, et al. Memory loss and memory reorganization patterns in temporal lobe epilepsy patients undergoing anterior temporal lobe resection, as demonstrated by pre-versus post-operative functional MRI. J Clin Neurosci. 2018; 55: 38–44.

Liu Z, Mikati M, Holmes GL. Mesial temporal sclerosis: pathogenesis and significance. Pediatr Neurol. 1995; 12: 5–16.

Long NM, Kahana MJ. Successful memory formation is driven by contextual encoding in the core memory network. Neuroimage. 2015; 119: 332–7.

Manns JR, Eichenbaum H. Evolution of declarative memory. Hippocampus. 2006; 16: 795808.

McLaren DG, Ries ML, Xu G, Johnson SC. A generalized form of context-dependent psychophysiological interactions (gPPI): a comparison to standard approaches. Neuroimage. 2012; 61: p. 1277–86.

McIntyre DC, Racine RJ. Kindling mechanisms: current progress on an experimental epilepsy model. Prog Neurobiol. 1986; 27: 1–12.

Meltzer JA, Constable RT. Activation of human hippocampal formation reflects success in both encoding and cued recall of paired associates. Neuroimage. 2005; 24: 384–97.

Milian ML, Zeltner M, Erb U, Klose K, Wagner L, Frings C, et al. Incipient preoperative reorganization processes of verbal memory functions in patients with left temporal lobe epilepsy. Epilepsy Behav. 2015; 42: 78–85.

Mottaghy FM, Shah NJ, Krause BJ, Schmidt D, Halsband U, Jancke L, et al. Neuronal correlates of encoding and retrieval in episodic memory during a paired-word association learning task: a functional magnetic resonance imaging study. Exp Brain Res. 1999; 128: 332–42.

Mueller SG, Laxer KD, Scanlon C, Garcia P, McMullen WJ, Loring DW, et al. Different structural correlates for verbal memory impairment in temporal lobe epilepsy with and without mesial temporal lobe sclerosis. Hum Brain Mapp. 2012; 33: 489–99.

Norman KA. How hippocampus and cortex contribute to recognition memory: revisiting the complementary learning systems model. Hippocampus. 2010; 20: 1217–27.

Oldfield RC. ‘The assessment and analysis of handedness: the Edinburgh inventory. Neuropsychologia. 1971; 9: 97–113.

Powell HW, Richardson MP, Symms MR, Boulby PA, Thompson PJ, Duncan JS, et al. Reorganization of verbal and nonverbal memory in temporal lobe epilepsy due to unilateral hippocampal sclerosis. Epilepsia. 2007; 48: 1512–25.

Richardson MP, Strange BA, Duncan JS, Dolan RJ. Preserved verbal memory function in left medial temporal pathology involves reorganisation of function to right medial temporal lobe. Neuroimage. 2003; 20: S112–9.

Richardson MP, Strange BA, Duncan JS, Dolan RJ. Memory fMRI in left hippocampal sclerosis: optimizing the approach to predicting postsurgical memory. Neurology. 2006; 66: p. 699–705.

Sadaghiani S, Esposito MD. Functional Characterization of the Cingulo-Opercular Network in the Maintenance of Tonic Alertness. Cereb Cortex. 2015; 25: 2763–73.

Saling MM. Verbal memory in mesial temporal lobe epilepsy: beyond material specificity. Brain. 2009; 132: 570–82.

Scoville WB, Milner B. Loss of recent memory after bilateral hippocampal lesions. J Neurol Neurosurg Psychiatry. 1957; 20: 11–21.

Sidhu MK, Stretton J, Winston GP, Bonelli S, Centeno M, Vollmar C, et al. A functional magnetic resonance imaging study mapping the episodic memory encoding network in temporal lobe epilepsy. Brain. 2013; 136: 1868–88.

Sidhu MK, Stretton J, Winston GP, McEvoy AW, Symms M, Thompson PJ, et al. Memory network plasticity after temporal lobe resection: a longitudinal functional imaging study. Brain. 2016; 139: 415–30.

Smith ML, Bigel M, Miller LA. Visual paired-associate learning: in search of material-specific effects in adult patients who have undergone temporal lobectomy. Epilepsy Behav. 2011; 20: 326–30.

Sperling MR, O’Connor MJ, Saykin AJ, Plummer C. Temporal lobectomy for refractory epilepsy. JAMA. 1996; 276: 470–5.

Squire LR, Zola-Morgan S. The medial temporal lobe memory system. Science. 1991; 253: 1380–6.

Tracy JI, Doucet GE. ‘Resting-state functional connectivity in epilepsy: growing relevance for clinical decision making. Curr Opin Neurol. 2015; 28: 158–65.

Tracy JI, Osipowicz KZ. A conceptual framework for interpreting neuroimaging studies of brain neuroplasticity and cognitive recovery. NeuroRehabilitation. 2011; 29: 331–8.

Vannest JK, Eaton P, Henkel D, Siegel M, Tsevat RK, Allendorfer JB, et al. Cortical correlates of self-generation in verbal paired associate learning. Brain Res. 2012; 1437: 104–14.

Wechsler D. WMS-IV: Wechsler memory scale-fourth edition. San Antonio TX: Pearson; 2009

Wechsler D. WAIS-IV: Administration and Scoring Manual. San Antonio TX: The Psychological Corporation; 2008.

Yassa MA, Stark CE. Pattern separation in the hippocampus. Trends Neurosci. 2011; 34: 515–25.

