## Supplementary material for "Computational Support, Not Primacy, Distinguishes Compensatory Memory Reorganization in Epilepsy": Suppliment_Material

#### Supplementary Methods

##### Section 1: Participant inclusion/exclusion criteria

All patients met the following criteria: unilateral temporal lobe seizure onset through surface video and EEG recordings; normal MRI or MRI evidence of pathology located within the epileptogenic mesial temporal lobe; and concordant PET finding of hypometabolism in the ictal temporal lobe. Patients were excluded from the study for any of the following reasons: previous brain surgery; extra-temporal/multifocal epilepsy; central nervous system illness other than epilepsy; contraindications to MRI; psychiatric diagnosis other than a Depressive Disorder or hospitalization for any disorder listed in the Diagnostic and Statistical Manual of Mental Disorders-V. Depressive Disorders were allowed given the high comorbidity of depression and epilepsy (Tracy *et al.*, 2007).

Patients were always assessed clinically before entering the scanner, and no patient was scanned if there was any sign or suggestion of a postictal state. As a policy, functional scans are cancelled and rescheduled if the patient or clinician reported a seizure the previous day or within a 24-hour period prior to the scan. Also, no patient either showed nor reported any signs of a seizure during the scanning procedure itself. All controls were recruited from the Thomas Jefferson University staff community, and were free of psychiatric or neurological disorders as determined by a health screening measure. All participants were right handed and native English speakers. The study was approved by the Institutional Review Board for Research with Human Subjects at Thomas Jefferson University. All participants provided a written informed consent.

#### Section 2: Paired-associate memory task design and details

The design and details of the three phases of the Paired-Associate Memory task are shown in Figures 1 and 2. During the word-pair encoding condition, the participant was required to silently read and memorize each randomized word pair, presented visually in the center of the screen. The pairs were presented sequentially for a duration of 4000ms each with a jitter of 800 – 1200ms inter-stimulus interval (average jitter 1000 msec.). Five word pairs were presented over a 25 second period. Following the fifth word pair, the participant was given a distraction math task involving addition and subtraction problems that lasted 25 seconds. Participants were required to identify if the viewed arithmetic problem was correct or incorrect, indicating their answer through a key press response. This distractor condition prevented covert rehearsal of the word pairs. In the final and third phase, the participant was given one of the words from the originally presented pair and then asked to recall (say aloud) the other member of the memorized pair. These single word cues were presented for 4000ms, each with a jitter of 800 – 1200ms inter-stimulus interval (average jitter 1000msec.). Five such word pair lists with distraction and recall phases were presented, with a total of 150 whole brain BOLD volumes acquired within the 375 seconds of the task.

#### Section 3: MRI data acquisition and pre-processing

To collect BOLD activity during the 6.25 minutes of PAM task. EPI images were acquired with the following parameters: 150 volumes, slices=33, parallel to AC-PC plane, TR=2.5s, TE=35ms, FOV=230mm, flip angle=90°, matrix=128*128 voxels, slice-thickness=4mm. Prior to the task, T1-MPRAGE images were acquired with the following parameters: 180 slices, 256*256 voxels; TR=640ms, TE=3.2ms, flip angle=8°, FOV=256mm, in-plane resolution 1mm$^2$ in positions identical to the functional scans for anatomical referencing.

*Preprocessing analyses:* FMRI data preprocessing for PAM task was performed on each subject using fMRIPrep 1.5.8 (Esteban *et al.*, 2019) based on Nipype tool 1.4.1 (Gorgolewski *et al.*, 2011).

Following the fMRIprep guideline, dicom images of T1 and task functional were converted into the standard Brain Imaging Data Structure (BIDS) format using dcm2niix (https://www.nitrc.org/projects/mricrogl/) program. To begin automated preprocessing steps, we first setup the installation of python (v 3.8.0) on PowerShell environment. Next, Docker Container (v 19.03) was utilized for running fMRIprep with docker-fmriprep command.

*Anatomical data preprocessing:* Each T1w (T1-weighted) volume was corrected for INU (intensity non-uniformity) using N4BiasFieldCorrection v2.1.0 (Tustison *et al.*, 2010) and skull-stripped using antsBrainExtraction.sh v2.1.0 (using the OASIS template). Spatial normalization to the ICBM 152 Nonlinear Asymmetrical template version 2009c [RRID: SCR_008796] was performed through nonlinear registration with the antsRegistration tool of ANTs v2.1.0 (Avants et al. 2008), [RRID: SCR_004757], using brain-extracted versions of both T1w volume and template. Brain tissue segmentation of cerebrospinal fluid (CSF), white-matter (WM) and gray-matter (GM) was performed on the brain-extracted T1w using fast (Zhang *et al.*, 2001), (FSL v5.0.9, RRID: SCR_002823).

*Functional data preprocessing:* Task PAM functional data of each participant was slice time corrected using 3dTshift from AFNI v16.2.07 (Cox and Hyde, 1997) and motion corrected using mcflirt (FSL v5.0.9) (Jenkinson *et al.*, 2002). "Fieldmap-less" distortion correction was performed by co-registering the functional image to the same-subject T1w image with intensity inverted (Wang *et al.*, 2017) constrained with an average field map template (Treiber *et al.*, 2016), implemented with antsRegistration (ANTs). This was followed by co-registration to the corresponding T1w using boundary-based registration (Greve and Fischl, 2009) with nine degrees of freedom, using flirt (FSL). Motion correcting transformations, field distortion correcting warp, BOLD-to-T1w transformation and T1w-to- template (MNI152NLin2009cAsym) warp were concatenated and applied in a single step using antsApplyTransforms (ANTs v2.1.0) using Lanczos interpolation. Physiological noise regressors were extracted applying CompCor (Behzadi *et al.*, 2007). Principal components were estimated for the two CompCor variants: temporal (tCompCor) and anatomical (aCompCor). A mask to exclude signal with cortical origin was obtained by eroding the brain mask, ensuring it only contained subcortical structures. Six tCompCor components were then calculated including only the top 5% variable voxels within that subcortical mask. For aCompCor, six components were calculated within the intersection of the subcortical mask and the union of CSF and WM masks calculated in T1w space, after their projection to the native space of each functional run. Frame-wise displacement (Power *et al.*, 2014) was calculated for each functional run using the implementation of Nipype.

The corresponding details of fMRIprep pipeline are available on documentation (see, https://fmriprep.readthedocs.io/en/latest/workflows.html). Finally, in SPM 12, the preprocessed normalized images were smoothed with an isotropic convolution Gaussian kernel of 8 mm (full-width half-maximum, FWHM). To maintain uniformities of results, participants, who had, a deviation of more than 3 mm or degree in max head motion were excluded from the analysis.

The preprocessed data were then entered in to the first level GLM analysis, utilizing 24 motion regressors (6 base motion parameters + 6 temporal derivatives of the 6 motion parameters + 12 quadratic terms of six motion parameters and their six temporal derivatives). Second-level analyses were performed to reveal the activation regions relevant to the PAM task key contrast of key interest (SME effect minus math control) using SPM12 (http://www.fil.ion.ucl.ac.uk/spm/software/spm12).

#### Section 4: Generalized psychophysiological interaction (gPPI) analyses

The processed GLM models of the PAM task were imported into the CONN toolbox to identify the differential connectivities with the seed that emerged during the SME as compared to the math control condition. At the subject-level, we generated a PPI regressor for each of the task conditions by calculating the element-by-element product between the psychological condition and the physiological regressor. We then estimated the whole brain functional connectivity (correlation strength) between the time-course of the ROI seed(s) and the gPPI regressor. The gPPI output (beta weights), reflecting the strength of the connection between the seed and the significant target areas in the brain were then converted to z-scores using the Fisher's z-transformation.

#### Section 5: Prior reports of memory reorganization in LTLE

The above picture of compensatory memory reorganization in LTLE is more complex than prior reports of reorganization would suggest. Most studies addressing reorganization have focused on activation change post-surgery (Cheung *et al.,* 2009). Those that have focused on pre-surgical status involving the use of a paired-associate verbal memory paradigm, have found increases in activation in the right hippocampus relative to healthy controls (Milian *et al.,* 2015), but have not reported the other regions we have highlighted here. More general verbal word list learning paradigms have found right-sided activations in TLE, mostly involving the hippocampus/parahippocampus (Dupont *et al.*, 2000; Powell *et al.,* 2007; Richardson *et al.*, 2003), or the frontal lobe (Lee *et al.*, 2008; Sidhu *et al.,* 2013), noting that we did not find reorganization effects, either in terms of activation magnitude or FC's, to involve the frontal lobe. Other studies of verbal memory have found intra-hemispheric shifts to the left inferior frontal or left superior temporal gyrus in LTLE (Dupont *et al.*, 2000), again differing from our results. In terms of FC, a host of studies in TLE have reported on FC's associated with the ictal hippocampus, with reductions in FC the most common finding (see review by Tracy and Doucet). Our data, however, is the first to make clear the importance of psychophysiological interactions between the primary computational hub (i.e., the hippocampus) and both regional and network involvements during actual memory processes. Yet, with regard to the extant literature in TLE, perhaps most important and novel about our study, is that none of the prior studies have examined reorganization patterns unique to intact, compensated memory performance.

#### Section 6: Methodologic and conceptual considerations relevant to our findings in LTLE

IQ and neuropsychological measures were only available on the TLE patients not the healthy controls. Thus, IQ differences could have existed between our groups. With this limitation in mind, we examined the association between IQ and our PAM performance measures. The result showed that they were not significantly correlated (r=.17). Out of this same concern we re-ran our SVM classifier of Intact versus Impaired status in LTLE, adding IQ and a baseline neuropsychological measure of verbal memory encoding (CVLT-II Total Learning). The six predictors marked in Table 5 remained significant. IQ was not a significant predictor (IQ, p-value=.48), however, the neuropsychological measure of verbal memory encoding did emerge as a significant classifier (CVLT-II Total Learning, AUC=.73, p-value<.05). This was expected, and suggests that the memory performance levels captured by our Intact/Impaired classification were not task specific and are consistent with the LTLE patients broader baseline memory skills. These supplementary data made clear that our SVM model is capturing activation magnitude and FC effects that cannot be accounted for by these cognitively-relevant baseline characteristics of our LTLE sample. While epilepsy duration was not a factor in our data, age of onset was, in that it bore some relation to the significant FC effects distinguishing Intact/Impaired patients. The implication of age of onset should be studied further, but our data points to the possibility that an earlier onset for LTLE lays down a set of increased connectivities with not just the ictal but also the non-ictal hippocampus. Interestingly, this finding bears some similarity to evidence in the domain of language, where early as opposed to late age of onset, appears to be associated with inter-hemispheric reorganization (Hertz-Pannier *et al.*, 2002).

We acknowledge that because cognitive reorganization requires both cortico-cortico and cortico-subcortical communication, no complete modelling of reorganization can occur without accounting for potential changes in white matter fibers that no doubt implement direct or indirect streams of information processing between many brain areas (De Benedictis and Duffau, 2011). Changes in such pathways may allow for network functionality and the maintenance of performance, even after removal of core computational areas such as the hippocampus. Thus, white matter connectivity is likely an important mediator of brain plasticity and cognitive reorganization, and we have not accounted for it here. Also, in this project, when referring to memory reorganization we are referring only to acquired and pathologically-driven reorganization, and not the natural remodeling of brain networks that ensures with learning, development, and aging (Duffau, 2008). We must also note that anti-epileptic medication can influence the blood oxygen level-dependent signal (Jansen *et al.*, 2006; Wandschneider *et al.*, 2017). However, the heterogeneity of the AED regimens (type, dose, etc.) in our sample precluded testing for AED effects. In light of the mixed distribution of medication within and across the TLE groups it seems unlikely that any of the Good/Poor Intact/Impaired performance differences we report can be traced to a specific medication. Also, the effect of medication would cancel out in any within TLE comparison. Importantly, we do not claim that the forms of compensatory reorganization we described for LTLE holds for all cognitive processes or tasks, nor for all forms of epilepsy. While we highlight several important hemisphere lateralized effects associated with Good/Poor and Intact/Impaired differences in LTLE, we emphasize that memory reorganization must be understood at the regional not the hemispheric level. Also, we realize that whenever there are shifts in hemispheric implementation of function, one must consider the possibility that the findings reflect a change in the normal inhibitory activity and control that goes on between the hemisphere, rather than a shift caused by pathology knocking out a function in a particular hemisphere. Unfortunately, our data provides no way to distinguish those possibilities, though we acknowledge our interpretation has generally assumed the latter. We acknowledge that there is no single way that the brain reorganizes, noting that any shifts or re-arrangements in the implementation of functions such as memory are happening in response to a large number of factors: the specific type of injury to brain gray and white matter, individual differences in learning history, age, gender, and the unique features of the underlying disease state. There has been an assumption in the TLE memory reorganization that non-dominant hemisphere findings are flawed substitutions for the injured dominant hemisphere, particularly when some of the non-dominant hemisphere effects set in late in the course of the disease, as evidenced by a relationship with early age of onset, a relationship we also find in our data. While their exact cognitive role in our LTLE sample is uncertain, it is clear that the regions we have revealed to be involved in reorganization are both compensatory and adaptive, not at all diminishing of performance. Lastly, we readily acknowledge that we did not explore the FC's at work with all the PAM task activations uniquely associated with Good/Poor or Intact/Compensated memory status in LTLE. Thus, while our data do demonstrate that certain changes in FC and communication are likely a necessary ingredient of successful reorganization, task mediated changes in communication with other regions are likely important as well. In a similar way, we must acknowledge that some TLE patients may have instituted regional changes in computational primacies or computational supports for memory processing elsewhere in the brain, but because such reorganization was individual and idiosyncratic, it failed to reach statistical significance in our group analyses.

#### Section 7: Data availability

The data that support the findings of this study are available within the article and its supplementary material. Additional data relevant to the study can be provided upon request from the corresponding author.

### Supplementary Table S1

| **MNI Coordinates of Significant Cluster Maxima for Subsequent Memory Effect minus Math Contrast Demonstrating Experimental Group Differences.** | | | | |
|---|---|---|---|---|
| | **Peak MNI Coordinates** | | | |
| **Region** | **k** | **x** | **y** | **z** |
| **LTLE minus HC** | | | | |
| R Parahippocampal gyrus | 508 | 14 | -24 | -18 |
| R Hippocampus | 371 | 32 | -4 | -14 |
| L Middle/superior frontal gyrus | 224 | -16 | 52 | 22 |
| R Middle/superior frontal gyrus | 112 | 4 | 38 | 12 |
| **RTLE minus HC** | | | | |
| L Superior parietal lobule | 427 | -26 | -68 | 36 |
| R Precentral gyrus | 168 | 46 | 8 | 24 |
| R Superior parietal lobule | 81 | 28 | -54 | 46 |
| L Fusiform gyrus | 57 | -30 | -78 | -8 |
| L Postcentral gyrus | 33 | -40 | -32 | 42 |
| The cluster thresholds correspond to corrected False Discovery Rate significance level of height $p$ <0.05. | | | | |

### References

Avants BB, Epstein CL, Grossman M, Gee JC. Symmetric diffeomorphic image registration with cross-correlation: evaluating automated labeling of elderly and neurodegenerative brain. Med Image Anal. 2008; 12: 26-41.

Behzadi Y, Restom K, Liau J, Liu TT. A component based noise correction method (CompCor) for BOLD and perfusion based Fmri. Neuroimage. 2007; 37: 90-101.

Cheung M, Chan AS, Lam JM, Chan YL. Pre- and postoperative fMRI and clinical memory performance in temporal lobe epilepsy. J Neurol Neurosurg Psychiatry. 2009; 80: 1099-106.

Cox RW, Hyde JS. Software tools for analysis and visualization of fMRI data. NMR Biomed. 1997; 10: 171-8.

De Benedictis A, Duffau H. Brain hodotopy: from esoteric concept to practical surgical applications. Neurosurgery. 2011; 68: 1709-23; discussion 23.

Duffau H. Brain plasticity and tumors. In: Pickard JD, Akalan N, Rocco C, Dolenc VV, Antunes JL, editors. Advances and Technical Standards in Neurosurgery. Springer Vienna; 2008. p. 3-33.

Dupont S, Van de Moortele PF, Samson S, Hasboun D, Poline JB, Adam C, et al. Episodic memory in left temporal lobe epilepsy: a functional MRI study. Brain. 2000; 123: 1722-32.

Esteban O, Markiewicz CJ, Blair RW, Moodie CA, Isik AI, Erramuzpe A, et al. fMRIPrep: a robust preprocessing pipeline for functional MRI. 2019; Nat Methods; 16: 111-16.

Gorgolewski K, Burns CD, Madison C, Clark D, Halchenko YO, Waskom ML, et al. Nipype: a flexible, lightweight and extensible neuroimaging data processing framework in python. Front Neuroinform. 2011; 5: 13.

Greve DN, Fischl B. Accurate and robust brain image alignment using boundary-based registration. Neuroimage. 2009; 48: 63-72.

Hertz-Pannier L. Chiron C, Jambaque I, Renaux-Kieffer V, Van de Moortele PF, Delalande O, et al. Late plasticity for language in a child's non-dominant hemisphere: a pre- and post-surgery fMRI study. Brain. 2002; 125: 361-72.

Jansen JFA, Aldenkamp AP, Marian HJM, Reijs RP, de Krom MC, Hofman PA, et al. Functional MRI reveals declined prefrontal cortex activation in patients with epilepsy on topiramate therapy. Epilepsy Behav. 2006; 9: 181-5.

Jenkinson MP, Brady BM, Smith S. Improved optimization for the robust and accurate linear registration and motion correction of brain images. Neuroimage. 2002; 17: 825-41.

Lee D, Swanson SJ, Sabsevitz DS, Hammeke TA, Winstanley FS, Possing ET, et al. Functional MRI and Wada studies in patients with interhemispheric dissociation of language functions. Epilepsy Behav. 2008; 13: 350-6.

Milian ML, Zeltner M, Erb U, Klose K, WagnerL, Frings C, et al. Incipient preoperative reorganization processes of verbal memory functions in patients with left temporal lobe epilepsy. Epilepsy Behav. 2015; 42: 78-85.

Powell HW, Richardson MP, Symms MR, Boulby PA, Thompson PJ, Duncan JS et al. Reorganization of verbal and nonverbal memory in temporal lobe epilepsy due to unilateral hippocampal sclerosis. Epilepsia. 2007; 48: 1512-25.

Power JD, Mitra A, Laumann TO, Snyder AZ, Schlaggar BL, Petersen SE. Methods to detect, characterize, and remove motion artifact in resting state fMRI. Neuroimage; 2014; 84: 320-41.

Richardson MP, Strange BA, Duncan JS, Dolan RJ. Preserved verbal memory function in left medial temporal pathology involves reorganisation of function to right medial temporal lobe. Neuroimage. 2003; 20: S112-9.

Sidhu MK, Stretton J, Winston GP, Bonelli S, Centeno M, Vollmar C, et al. A functional magnetic resonance imaging study mapping the episodic memory encoding network in temporal lobe epilepsy. Brain. 2013; 136: 1868-88.

Tracy JI, Lippincott C, Mahmood T, Waldron B, Kanauss K, Glosser D, et al. Are depression and cognitive performance related in temporal lobe epilepsy? Epilepsia. 2007; 48: 2327-35.

Treiber JM, White NS, Steed TC, Bartsch H, Holland D, Farid N, et al. Characterization and Correction of Geometric Distortions in 814 Diffusion Weighted Images. PLoS One. 2016; 11: e0152472.

Tustison NJ, Avants BB, Cook PA, Zheng Y, Egan A, Yushkevich PA, et al. N4ITK: improved N3 bias correction. IEEE Trans Med Imaging. 2010; 29: 1310-20.

Wandschneider B, Burdett J, Townsend L, Hill A, Thompson PJ, Duncan JS, et al. Effect of topiramate and zonisamide on fMRI cognitive networks. Neurology. 2017; 88: 1165-71.

Wang S, Peterson DJ, Gatenby JC, Li W, Grabowski TJ, Madhyastha TM. Evaluation of Field Map and Nonlinear Registration Methods for Correction of Susceptibility Artifacts in Diffusion MRI. Front Neuroinform. 2017; 11: 17.

Zhang Y, Brady M, Smith S. Segmentation of brain MR images through a hidden Markov random field model and the expectation-maximization algorithm. IEEE Trans Med Imaging. 2001; 20: 45-57.

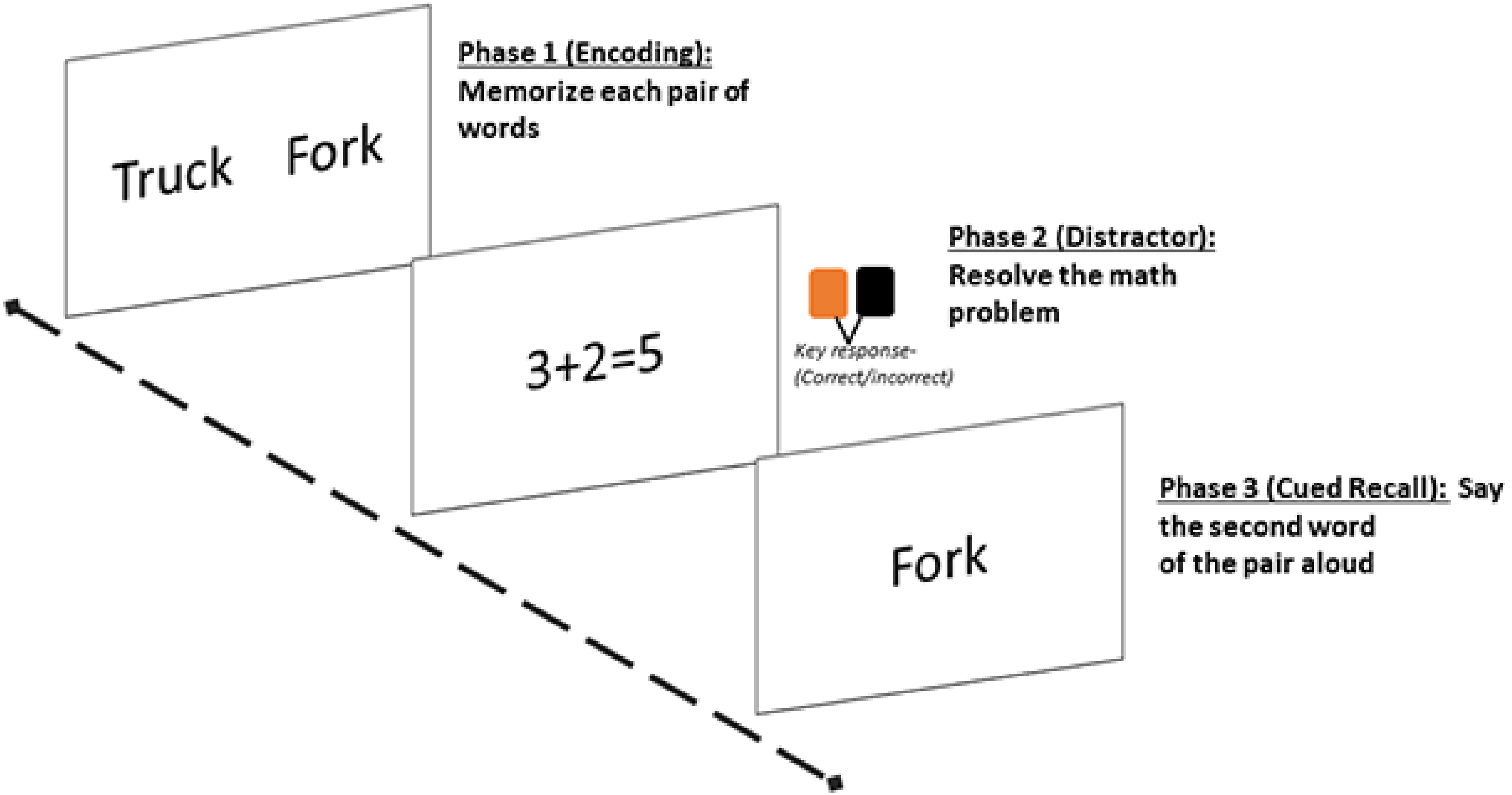

Supplement Section, Figure 1: The fMRI paired-associate verbal learning task and memory task comprised three phases. The first phase exhibited pairs of words. The participant was instructed to silently read and memorize each pair. The second phase, a distractor condition, presented simple math problems, with the participant instructed to press a key indicating if the answer listed was correct or not. In the third phase, a single word from the target pairs was presented centrally, with the participant instructed to say aloud the other member of the pair (cued recall).

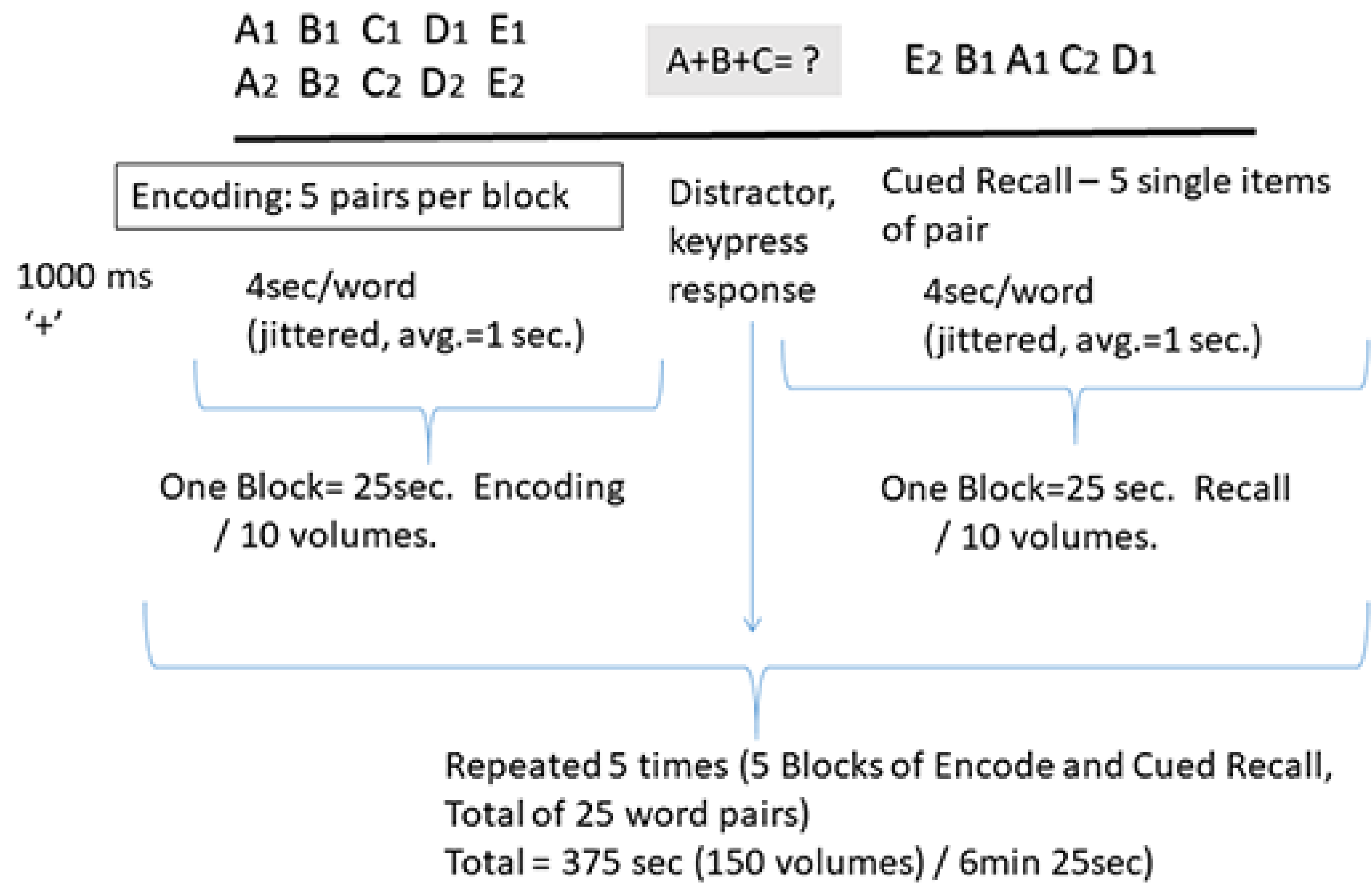

Supplement Section., Figure 2: Detailed schematic of events in the paired-associate verbal learning memory paradigm. Following a 1-second fixation, participants were administered 5 blocks, each block presenting 5 word pairs during an encoding phase. Next, intervening math problems served as a distractor condition. Lastly, a cued recall phase was used to collect responses and determine forgotten versus successfully encoded pairs, allowing for calculation of the subsequent memory and forgetting effects. This cycle was repeated 5 times, through 5 distinct, randomized word-pair lists. The contrast of interest compared the encoding phase (SME, subsequent memory effect) to the math distractor phase. The total time, 375 seconds, was time required administer the task and acquire 150 whole brain BOLD volumes.

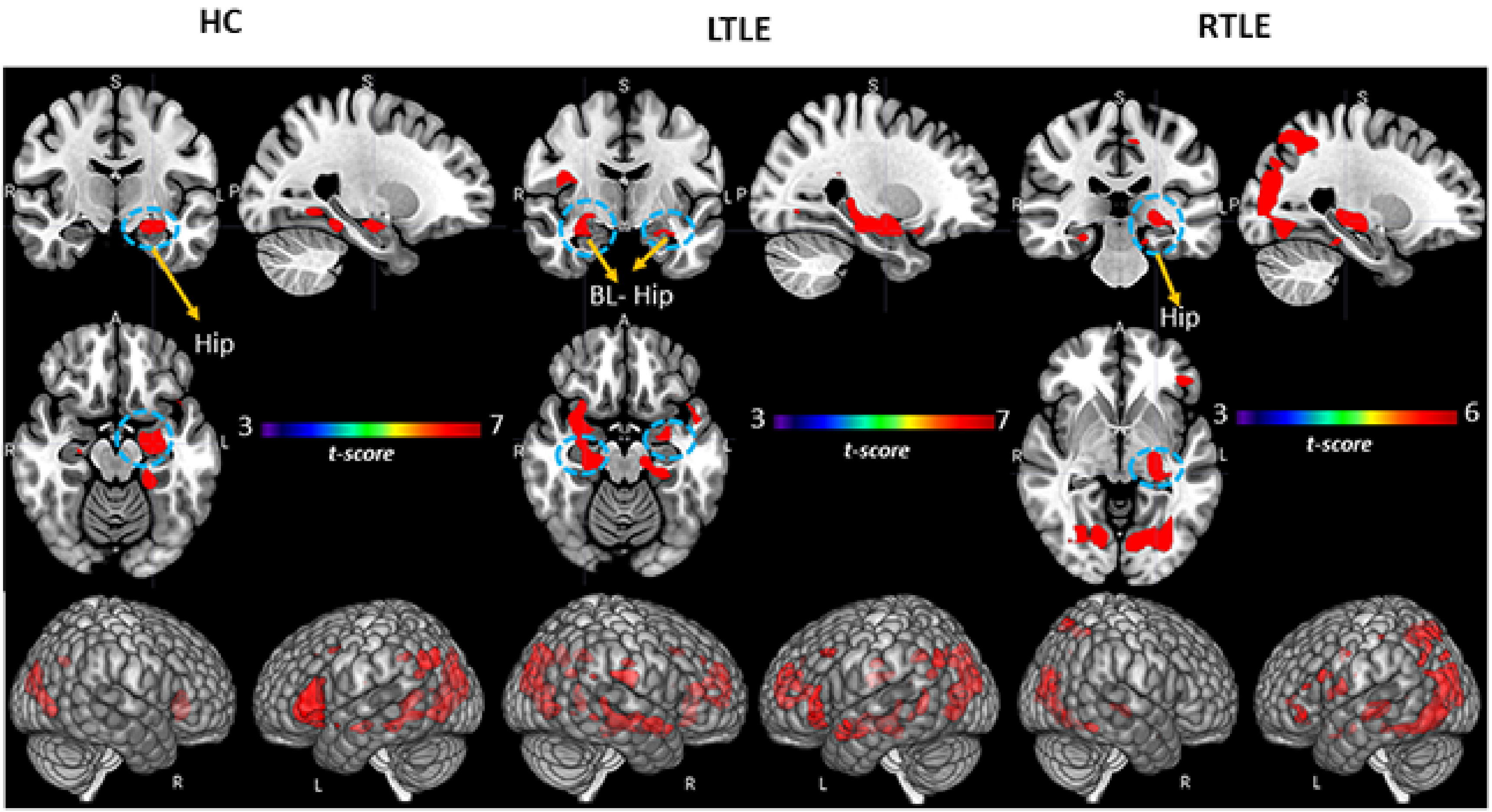

Supplement Section, Figure 3: Regional activation associated with subsequent memory effect (SME) minus math condition contrast. Surface rendering and slices in vertical panels show significant activation for HC, LTLE, and RTLE groups (*pFDR* corrected<.01, cluster level). BL: Bilateral; Hip: Hippocampus
